# RNA methyltransferase CMTR-1 inhibition activates a GATA transcription factor-mediated protective immune response

**DOI:** 10.64898/2025.12.09.693339

**Authors:** Annesha Ghosh, Jogender Singh

## Abstract

Organisms can activate innate immunity not only by detecting pathogens but also by sensing disturbances to essential cellular processes. Such damage-based surveillance provides a means to identify cellular disruptions that prime host immunity even in the absence of infection. To uncover pathways that enhance immunity, we perform a forward genetic screen in *Caenorhabditis elegans* for mutants with elevated intestinal immune activation. This screen identifies a hypomorphic allele of the conserved mRNA cap 2’-O-methyltransferase *cmtr-1*, which strongly induces immune effector expression. Transcriptomic profiling reveals the broad activation of innate immune responses, and functional analyses demonstrate that this response requires the GATA transcription factor ELT-2. Notably, loss of *cmtr-1* increases resistance to *Pseudomonas aeruginosa*, indicating that reduced RNA cap methylation can elicit protective immunity. Together, these findings reveal a previously unrecognized mechanism of immune surveillance and demonstrate that perturbation of CMTR-1 serves as a danger signal, triggering an ELT-2-dependent protective immune response.

## Introduction

Animals have evolved diverse strategies to detect and respond to microbial pathogens (Blander & Sander, 2012; Medzhitov, 2007; Singh & Aballay, 2020). A major mode of pathogen detection involves the recognition of pathogen-associated molecular patterns by conserved pattern-recognition receptors, which initiate signaling cascades that activate innate immunity (Janeway & Medzhitov, 2002; Medzhitov, 2007). In addition to direct pathogen sensing, hosts have also developed mechanisms to detect damage to endogenous cellular components caused by infection or stress. Such damage-associated molecular patterns and stress-response pathways provide an additional and often pathogen-independent means of immune activation. In *Caenorhabditis elegans* and other organisms, disruptions to core cellular processes, including protein translation, endoplasmic reticulum homeostasis, mitochondrial function, and proteasomal activity, trigger innate immune responses even in the absence of pathogen infection (Bettigole & Glimcher, 2015; Dunbar *et al*, 2012; Ghosh & Singh, 2024; Grover *et al*, 2024; McEwan *et al*, 2012; Melo & Ruvkun, 2012; Pellegrino *et al*, 2014; Rao *et al*, 2025; Ghosh & Singh, 2026). This capacity to sense cellular dysfunction as a danger signal allows hosts to respond to a broad spectrum of pathogens without relying exclusively on specific microbial ligands. Moreover, because cellular damage pathways can be activated independently of infection, they offer a conceptual framework for uncovering perturbations that elicit protective immune states through mild, hormetic activation of stress and immune signaling.

Mild activation or priming of innate immunity can enhance host resistance to a variety of microbes. In *C. elegans*, for instance, neural circuits exert tonic inhibitory control over intestinal immunity (Singh & Aballay, 2020). Genetic or functional disruption of specific neurons, neuronal receptors, or neuropeptide pathways leads to elevated immune gene expression and increased survival upon infection with diverse bacterial pathogens (Cao & Aballay, 2016; Kawli & Tan, 2008; Styer *et al*, 2008; Sun *et al*, 2011; Yu *et al*, 2018). These and other findings illustrate that protective immune responses can arise from perturbations that do not involve pathogen exposure. Given that hosts can mount immune responses by detecting intrinsic cellular stress, systematic identification of such perturbations may uncover endogenous pathways that enhance pathogen resistance. This concept raises the possibility that controlled interference with core cellular processes could be exploited to identify mechanisms that enhance host defense.

In this study, we used a forward genetic screen in a *clec-60p::gfp* reporter background to identify *C. elegans* mutants with elevated basal immune activation. Our screen uncovered a hypomorphic allele of the conserved mRNA cap methyltransferase CMTR-1, which robustly activated *clec-60* expression. Transcriptomic profiling revealed that the isolated *cmtr-1(jsn21)* mutant upregulates a broad set of innate immune genes, and functional assays demonstrated that this immune activation enhances resistance to *Pseudomonas aeruginosa* infection. A transcription factor RNA interference (RNAi) library screen, combined with bioinformatic analyses, identified the intestinal GATA transcription factor ELT-2 as the key regulator of the immune program induced by *cmtr-1* loss-of-function. Together, these findings demonstrated that perturbation of CMTR-1, a conserved regulator of mRNA cap modification, serves as a danger signal that triggers a protective immune response. Our results further illustrate how disruption of essential cellular processes can be leveraged to uncover mechanisms that enhance host defense against pathogenic microbes.

## Results

### A recessive mutation in the mRNA cap methyl transferase CMTR-1 increases *clec-60p::gfp* levels

To identify mutants with enhanced immune activity, we performed a forward genetic screen using the *C. elegans* reporter strain *clec-60p::gfp*. This reporter monitors expression of the C-type lectin *clec-60*, an innate immune effector induced by diverse pathogens (Irazoqui *et al*, 2010). The screen was designed to isolate mutants exhibiting increased *clec-60p::gfp* expression, with the expectation that such mutants would possess augmented immunity against pathogen challenge (Fig 1A). Most of the isolated mutants were either sterile or failed to reach adulthood and died prematurely, consistent with previous findings that sterile mutants show heightened immune activation and that excessive immune activity impairs development (Cheesman *et al*, 2016; Miyata *et al*, 2008). Nonetheless, we identified a fertile and viable mutant with markedly elevated *clec-60p::gfp* expression (Fig 1B, 1C).

**Figure 1:**
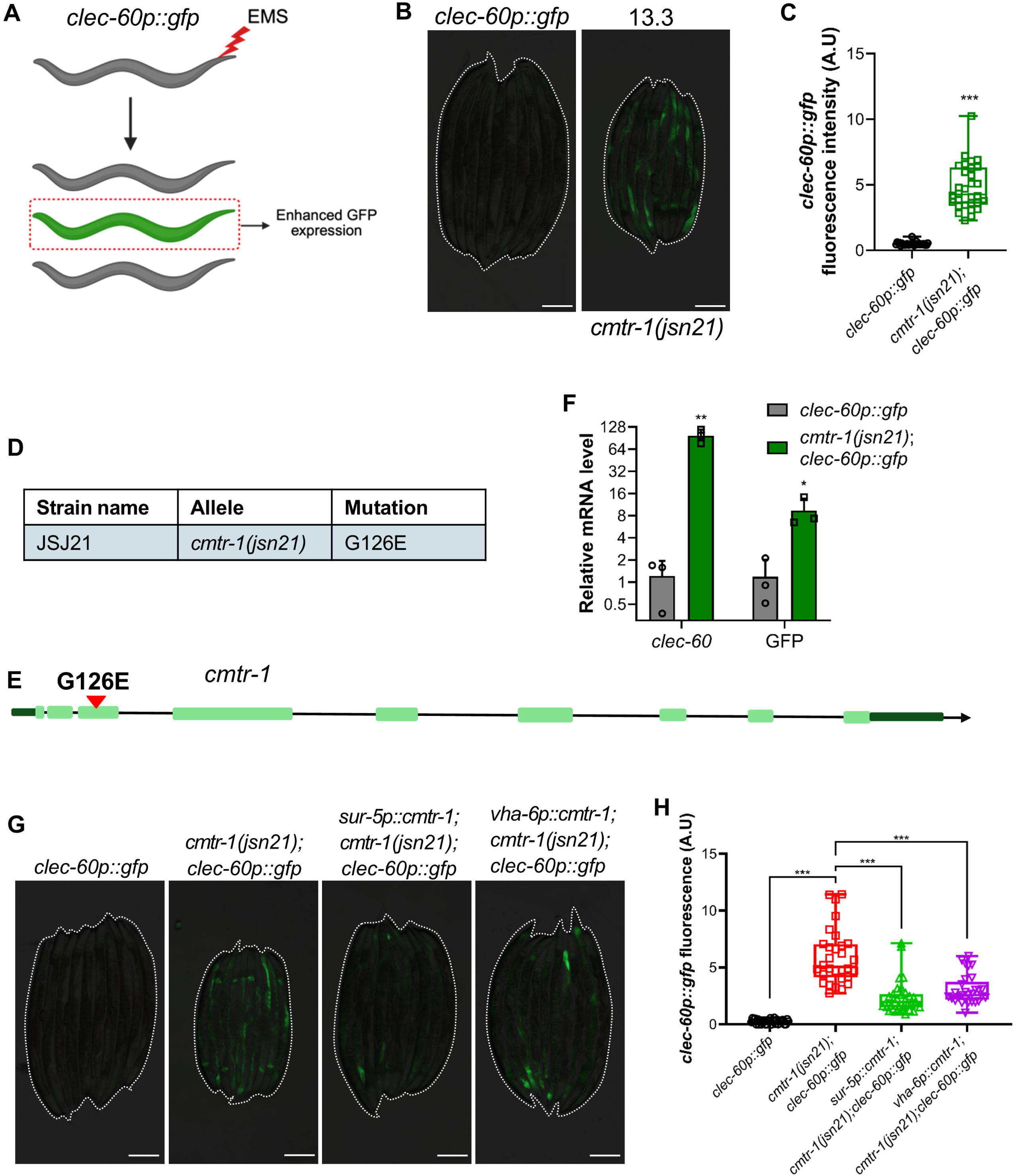
A recessive mutation in the RNA cap methyltransferase CMTR-1 increases *clec-60p::gfp* levels. (A) Scheme for a forward genetic screen for mutants that have an enhanced GFP in *clec-60p::gfp* worms. The worm illustrations were created using <u>BioRender</u>. (B) Representative fluorescence images of the wild-type parental strain *clec-60p::gfp* and the *cmtr-1(jsn21)* mutant isolated from the forward genetic screen with enhanced GFP levels. Dotted outlines indicate worm positions. Scale bar = 200 μm. (C) Quantification of GFP levels of *clec-60p::gfp* and *cmtr-1(jsn21);clec-60p::gfp* worms. \*\*\**p*< 0.001 via t-test (*n* = 30 worms each). (D) Summary of the *cmtr-1(jsn21)* allele. (E) Mapping of the *cmtr-1(jsn21)* allele identified in the forward genetic screen. (F) Quantitative reverse transcription-PCR (qRT-PCR) analysis of the immune gene *clec-60* and the transgene *gfp* in *clec-60p::gfp* and *cmtr-1(jsn21);clec-60p::gfp* worms. \**p*<0.05 and \*\**p*< 0.01 via t-test. Data represent the mean and standard deviation from three independent experiments. (G) Representative fluorescence images of *clec-60p::gfp*, *cmtr-1(jsn21);clec-60p::gfp*, *sur-5p::cmtr-1;cmtr-1(jsn21);clec-60p::gfp*, and *vha-6p::cmtr-1;cmtr-1(jsn21);clec-60p::gfp* worms. Dotted outlines indicate worm positions. Scale bar = 200 μm. (H) Quantification of GFP levels of *clec-60p::gfp*, *cmtr-1(jsn21);clec-60p::gfp*, *sur-5p::cmtr-1;cmtr-1(jsn21);clec-60p::gfp*, and *vha-6p::cmtr-1;cmtr-1(jsn21);clec-60p::gfp* worms. \*\*\**p*< 0.001 via t-test (*n* = 29-31 worms each).

This mutant was backcrossed six times to the parental strain, followed by whole-genome sequencing. Analysis of single-nucleotide polymorphisms and RNAi-based confirmation identified *cmtr-1* as the causal gene responsible for increased *clec-60p::gfp* expression (Fig S1A, S1B). *cmtr-1* encodes a methyltransferase that catalyzes the ribose 2’-O-methylation of the first transcribed nucleotide (Bélanger *et al*, 2010). The isolated allele, *cmtr-1(jsn21)*, contained a single G → A substitution leading to a Glycine (G) to Glutamic acid (E) substitution at residue 126 (Fig 1D, 1E). This mutation lies within the G-patch domain of CMTR-1, which mediates interaction with the RNA helicase DDX-15 (Toczydlowska-Socha *et al*, 2018; Inesta-Vaquera *et al*, 2018; Meisel *et al*, 2024). Previous studies have shown that this interaction can modulate CMTR-1 methyltransferase activity, either positively or negatively, depending on the context (Toczydlowska-Socha *et al*, 2018; Inesta-Vaquera *et al*, 2018). We therefore examined the inheritance pattern of *cmtr-1(jsn21)* with respect to *clec-60p::gfp* upregulation and found that the phenotype segregates recessively (Fig S1C), suggesting that the mutation likely reduces CMTR-1 activity to induce *clec-60* expression.

To confirm that *cmtr-1(jsn21)* worms indeed had increased activity of the *clec-60* promoter, we measured mRNA levels for the gene encoding green fluorescent protein (GFP). Indeed, *cmtr-1(jsn21);clec-60p::gfp* worms exhibited significantly increased GFP mRNA compared with *clec-60p::gfp* controls (Fig 1F). Consistent with reporter induction, *clec-60* mRNA levels were also upregulated in *cmtr-1(jsn21)* mutants, confirming that *cmtr-1* loss enhances *clec-60* expression at the transcriptional level. To establish that loss of *cmtr-1* function caused this phenotype, we expressed wild-type *cmtr-1* under the pan-tissue *sur-5* promoter. Because CMTR-1 is broadly expressed in *C. elegans* (Meisel *et al*, 2024), we first attempted pan-tissue rescue and observed near-complete suppression of elevated *clec-60p::gfp* expression (Fig 1G, 1H). Given that *clec-60* is predominantly expressed in the intestine, we next tested whether intestinal expression of *cmtr-1* was sufficient to rescue the phenotype. Expression of wild-type *cmtr-1* from the intestine-specific *vha-6* promoter significantly rescued *clec-60p::gfp* upregulation, demonstrating that intestinal CMTR-1 is sufficient to regulate *clec-60* expression (Fig 1G, H).

### Loss of *cmtr-1* enhances resistance to *Pseudomonas aeruginosa*

We next tested whether elevated *clec-60p::gfp* expression in *cmtr-1(jsn21)* mutants correlated with protection against *P. aeruginosa* PA14 infection. *P. aeruginosa* PA14 employs multiple mechanisms to kill *C. elegans* (Tan *et al*, 1999). When PA14 is cultured on minimal medium, infection-mediated lethality develops gradually over several days and is therefore termed “slow killing.” In contrast, growth of PA14 on high-osmolarity medium results in rapid lethality within a few hours, referred to as “fast killing.” We found that *cmtr-1(jsn21)* mutants exhibited significantly enhanced resistance to *P. aeruginosa* infection under both fast-killing (Fig S2A) and slow-killing conditions (Fig 2A). For the remainder of the study, we focused on the slow-killing model to investigate *P. aeruginosa* pathogenesis. Consistent with the mutant phenotype, RNAi-mediated knockdown of *cmtr-1* also increased survival on *P. aeruginosa* relative to the empty vector control (Fig S2B). Moreover, expression of *cmtr-1* under either pan-tissue or intestine-specific promoters suppressed the enhanced survival phenotype, demonstrating that improved pathogen resistance arises specifically from loss of *cmtr-1* function (Fig 2B, 2C).

**Figure 2:**
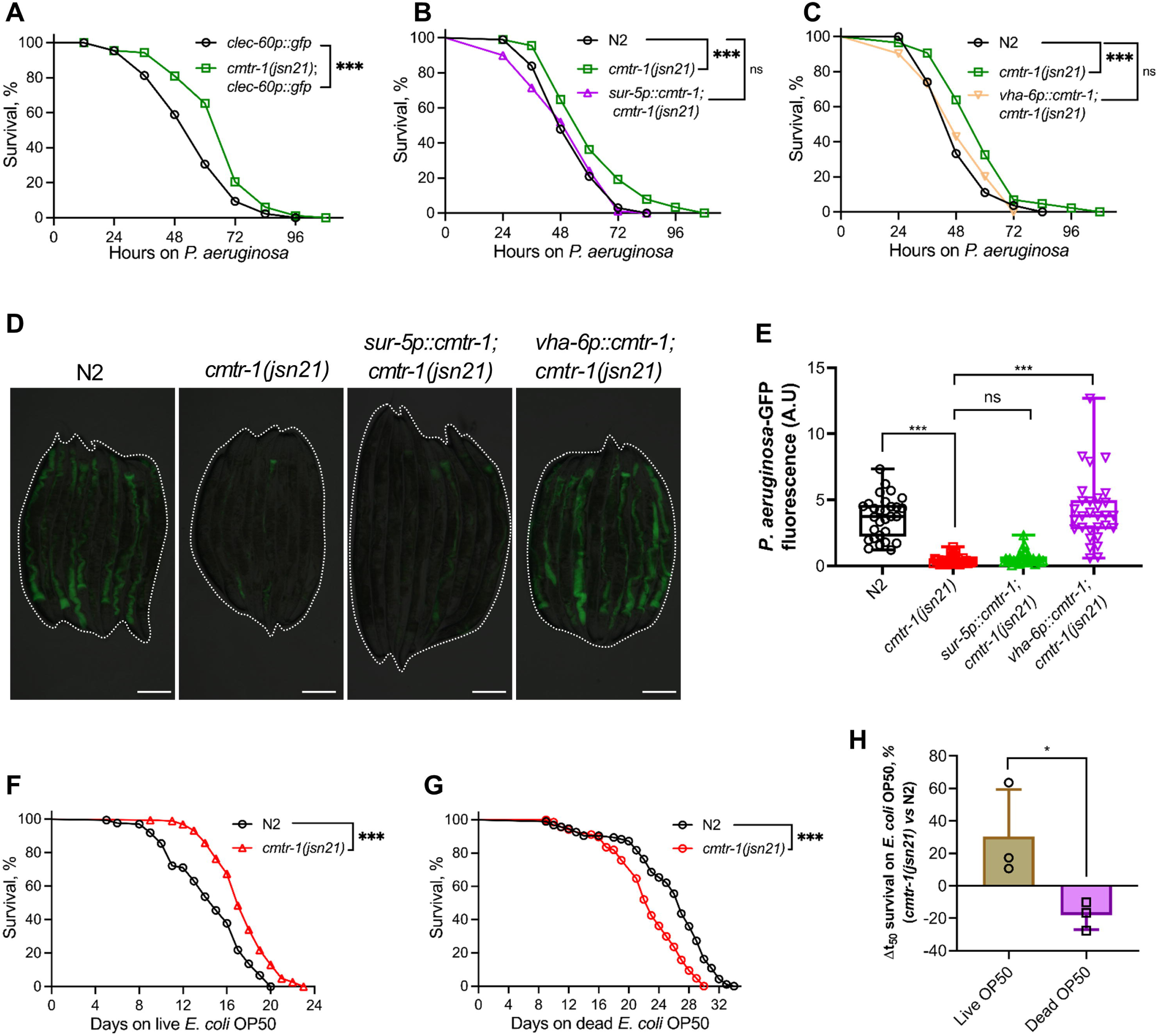
Loss of *cmtr-1* enhances resistance to *P. aeruginosa*. (A) Representative survival plots of *clec-60p::gfp* and *cmtr-1(jsn21);clec-60p::gfp* worms on *P. aeruginosa* PA14 at 25°C. \*\*\**p*< 0.001 for the mutant as compared to the control worms (*n* = 90 per condition). (B) Representative survival plots of N2, *cmtr-1(jsn21)*, and *sur-5p::cmtr-1;cmtr-1(jsn21)* worms on *P. aeruginosa* PA14 at 25°C. \*\*\**p*< 0.001 between N2 and *cmtr-1(jsn21)*, and non-significant between N2 and *sur-5p::cmtr-1;cmtr-1(jsn21)* (*n* = 90 per condition). (C) Representative survival plots of N2, *cmtr-1(jsn21)*, and *vha-6p::cmtr-1;cmtr-1(jsn21)* worms on *P. aeruginosa* PA14 at 25°C. \*\*\**p*< 0.001 between N2 and *cmtr-1(jsn21)*, and non-significant between N2 and *vha-6p::cmtr-1;cmtr-1(jsn21)* (*n* = 70 for N2, 86 for *cmtr-1(jsn21)*, and 114 for *vha-6p::cmtr-1;cmtr-1(jsn21)*). (D) Representative fluorescence images of N2, *cmtr-1(jsn21)*, *sur-5p::cmtr-1;cmtr-1(jsn21)*, and *vha-6p::cmtr-1;cmtr-1(jsn21)* worms incubated on *P. aeruginosa*-GFP for 24 hours at 25°C. Dotted outlines indicate worm positions. Scale bar = 200 μm. (E) Quantification of GFP levels of N2, *cmtr-1(jsn21)*, *sur-5p::cmtr-1;cmtr-1(jsn21)*, and *vha-6p::cmtr-1;cmtr-1(jsn21)* worms incubated on *P. aeruginosa*-GFP for 24 hours at 25°C. \*\*\**p*< 0.001 and non-significant (ns) via t-test (*n* = 29-30 worms each). (F) Representative survival curves of N2 and *cmtr-1(jsn21)* worms fed on live *E. coli* OP50. \*\*\**p*< 0.001 for *cmtr-1(jsn21)* compared to N2 (*n* = 173 for N2 and 193 for *cmtr-1(jsn21)*). (G) Representative survival curves of N2 and *cmtr-1(jsn21)* worms fed on kanamycin-killed *E. coli* OP50. \*\*\**p*< 0.001 for *cmtr-1(jsn21)* compared to N2 (*n* = 97 for N2 and 69 for *cmtr-1(jsn21)*). (H) The percentage change in mean survival of *cmtr-1(jsn21)* worms relative to N2 worms on live and kanamycin-killed *E. coli* OP50. \**p*< 0.05 via the t-test. Data represent the mean and standard deviation from three independent experiments.

Because intestinal colonization by *P. aeruginosa* strongly influences survival under slow-killing conditions (Das *et al*, 2024; Tan *et al*, 1999), we next examined bacterial accumulation in the gut. *cmtr-1(jsn21)* mutants showed markedly reduced intestinal colonization (Fig 2D, 2E). Notably, intestine-specific expression of *cmtr-1* restored intestinal colonization, whereas pan-tissue expression failed to do so (Fig 2D, 2E). This observation was unexpected because both rescue constructs effectively suppressed the enhanced survival phenotype. One possible explanation is that the expression pattern driven by the *sur-5* promoter does not fully recapitulate endogenous *cmtr-1* expression. Additionally, previous studies have demonstrated that reduced bacterial colonization and enhanced survival are not always directly correlated (Ghosh & Singh, 2024; Rao *et al*, 2025; Peterson *et al*, 2022). Nevertheless, these findings established that intestinal CMTR-1 is sufficient to restore colonization in *cmtr-1(jsn21)* mutants. Collectively, these results demonstrated that loss of *cmtr-1* enhances immune activity, leading to reduced *P. aeruginosa* colonization and improved host survival.

Because increased survival on a pathogen can reflect broader lifespan extension (Naim *et al*, 2021; Xia *et al*, 2019; Soo *et al*, 2023), we examined the lifespan of *cmtr-1(jsn21)* mutants on *E. coli* OP50 in the presence of 5-fluorodeoxyuridine (FUdR). Intriguingly, *cmtr-1(jsn21)* worms exhibited extended lifespan on *E. coli* OP50 (Fig 2F). To test whether this resulted from the mild pathogenicity of *E. coli* OP50 (Sánchez-Blanco & Kim, 2011), we measured lifespan on kanamycin-killed *E. coli* OP50. Under these conditions, *cmtr-1(jsn21)* mutants exhibited a significantly reduced lifespan relative to wild-type worms in the presence of FUdR (Fig 2G, 2H). Because FUdR can influence lifespan in a genotype-dependent manner (Van Raamsdonk & Hekimi, 2011), we further examined longevity in the absence of FUdR. Under these conditions, *cmtr-1(jsn21)* mutants displayed a lifespan comparable to that of wild-type worms when maintained on live bacteria (Fig S2C, S2D), but exhibited significantly reduced survival on dead *E. coli* OP50 (Fig S2E, S2F). Together, these findings indicate that although *cmtr-1(jsn21)* mutants exhibit enhanced immune responses and increased resistance to bacterial infection, they do not possess an intrinsically extended lifespan.

### *cmtr-1(jsn21)* mutants upregulate innate immune genes and downregulate translation-related genes

To characterize genome-wide transcriptional changes associated with *cmtr-1* loss, we performed RNA sequencing of *cmtr-1(jsn21)* mutants. Relative to wild-type worms, mutants showed 1,759 differentially expressed genes, with 1,217 significantly upregulated and 542 significantly downregulated (Fig 3A; Table S1). Gene Ontology (GO) analysis revealed strong enrichment of innate immune response genes among upregulated transcripts (Fig 3B; Fig S3A), whereas downregulated genes were enriched for components of the translation machinery (Fig 3C; Fig S3B).

**Figure 3:**
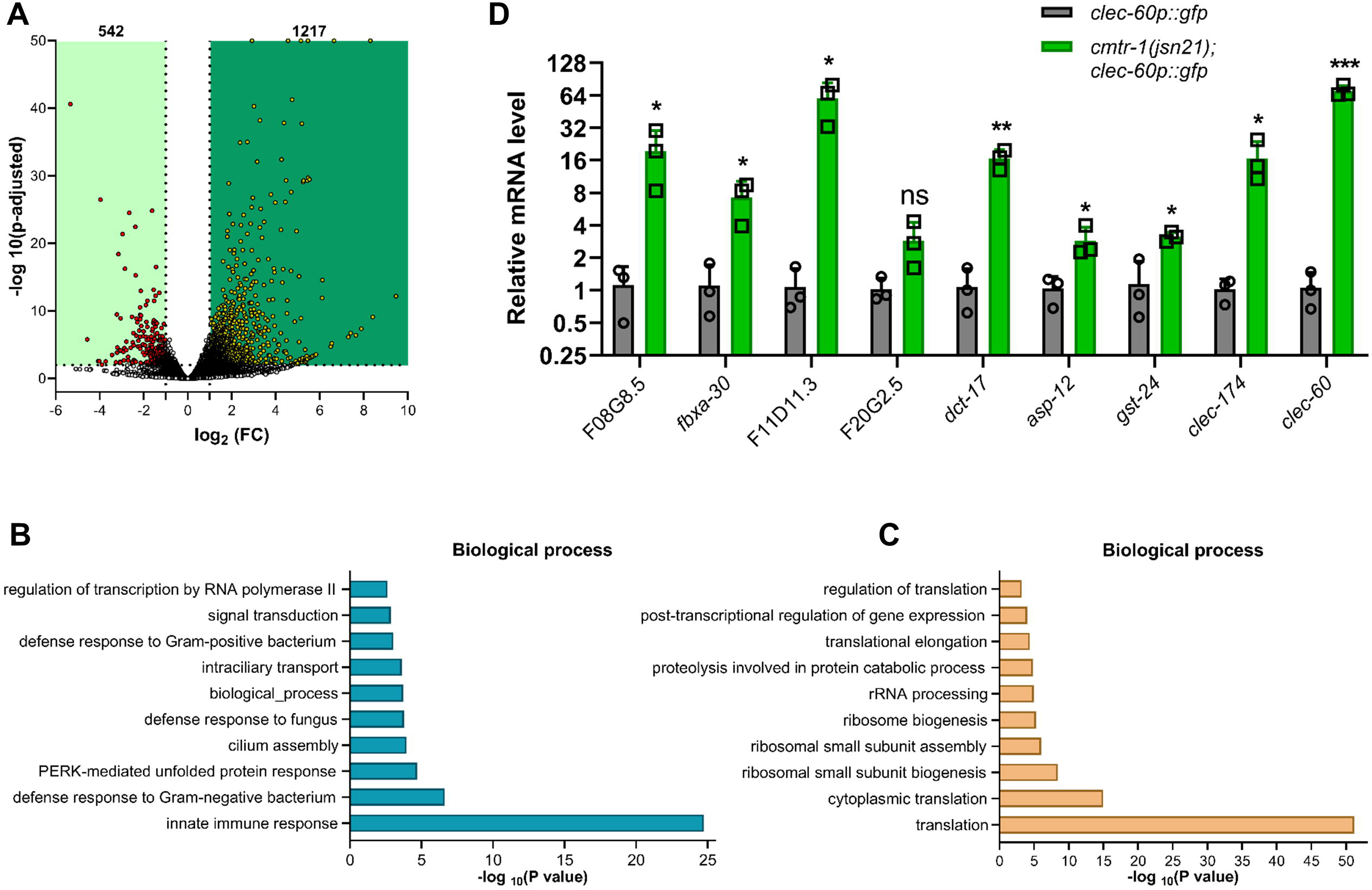
*cmtr-1(jsn21)* mutants upregulate innate immune genes and downregulate translation-related genes. (A) Volcano plot of upregulated and downregulated genes in *cmtr-1(jsn21);clec-60p::gfp* worms compared to control *clec-60p::gfp* worms grown on *E. coli* OP50. (B)-(C) Gene Ontology (GO) enrichment analysis for biological processes for upregulated (B) and downregulated (C) genes in *cmtr-1(jsn21);clec-60p::gfp* worms compared to control *clec-60p::gfp* worms grown on *E. coli* OP50. (D) qRT-PCR analysis of innate immune response genes in *clec-60p::gfp* and *cmtr-1(jsn21);clec-60p::gfp* worms grown on *E. coli* OP50. \*\*\**p*< 0.001, \*\**p*< 0.01, \**p*< 0.05, and ns, non-significant via t-test. Data represent the mean and standard deviation from three independent experiments.

To validate these findings, we quantified mRNA levels of selected immune genes by quantitative reverse transcription-PCR (qRT-PCR). *cmtr-1(jsn21)* mutants exhibited significantly elevated expression of these genes relative to wild-type worms (Fig 3D), supporting the RNA sequencing results. Together, these analyses demonstrated that loss of *cmtr-1* activates a broad innate immune transcriptional program.

### Enhanced immunity in *cmtr-1(jsn21)* is independent of major innate immune pathways

Because *cmtr-1(jsn21)* mutants exhibited reduced expression of translation-related genes (Fig 3C), we next investigated whether immune activation in these animals resulted from impaired protein synthesis. Inhibition of translation initiation is known to activate innate immune responses and enhance resistance to *P. aeruginosa* under slow-killing conditions (Ghosh & Singh, 2024). Since *cmtr-1(jsn21)* worms also displayed reduced expression of translation initiation factors (Table S1), we asked whether their increased survival on *P. aeruginosa* was linked to reduced translation. To test this, we knocked down the translation initiation factors *eif-2α* and *ifg-1* in wild-type and *cmtr-1(jsn21)* worms and examined survival during *P. aeruginosa* infection. Consistent with previous findings (Ghosh & Singh, 2024), inhibition of *eif-2α* and *ifg-1* increased the survival of wild-type worms (Fig S4A). However, knockdown of either factor did not further enhance the survival of *cmtr-1(jsn21)* mutants (Fig S4B), suggesting that the increased pathogen resistance of *cmtr-1(jsn21)* worms may be associated with reduced translation.

Translation inhibition has been shown to activate immune responses via the bZIP transcription factor ZIP-2, thereby enhancing resistance to *P. aeruginosa* (Dunbar *et al*, 2012; Ghosh & Singh, 2024; McEwan *et al*, 2012). We therefore examined whether ZIP-2 was required for the enhanced pathogen resistance observed in *cmtr-1(jsn21)* mutants. However, the strong upregulation of *clec-60p::gfp* in *zip-2(tm4248);cmtr-1(jsn21)* worms indicated that the *cmtr-1(jsn21)* phenotype does not depend on ZIP-2 (Fig 4A, 4B). Consistently, ZIP-2 was also dispensable for the reduced intestinal colonization (Fig. 4C, 4D) and enhanced survival during infection observed in *cmtr-1(jsn21)* mutants (Fig. 4E, 4F). Together, these findings demonstrated that immune activation in *cmtr-1(jsn21)* worms occurs independently of the canonical ZIP-2-mediated translation surveillance pathway.

**Figure 4:**
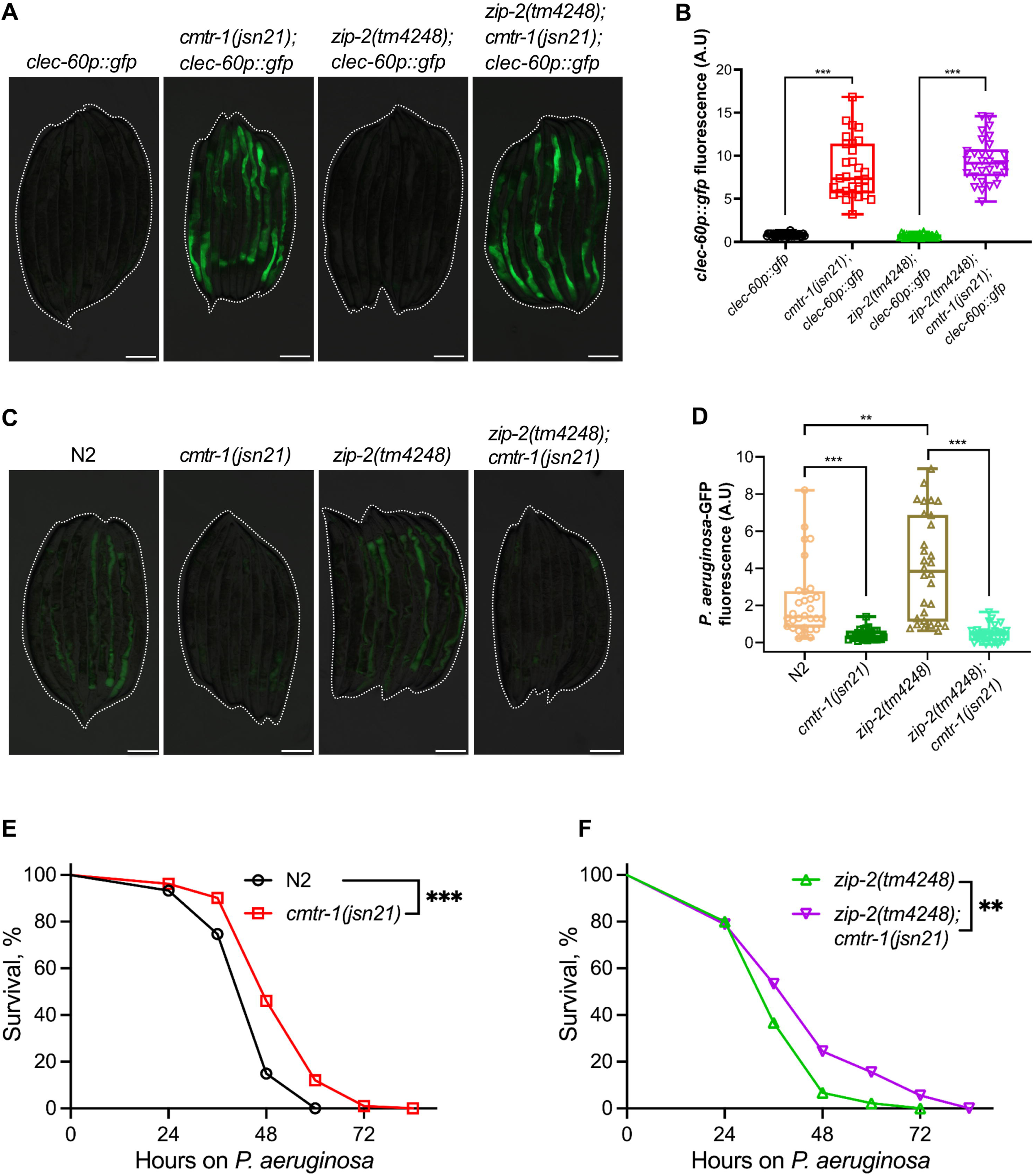
Enhanced immunity in *cmtr-1(jsn21)* worms is independent of ZIP-2. (A) Representative fluorescence images of *clec-60p::gfp*, *cmtr-1(jsn21);clec-60p::gfp*, *zip-2(tm4248);clec-60p::gfp*, and *zip-2(tm4248);cmtr-1(jsn21);clec-60p::gfp* worms. Dotted outlines indicate worm positions. Scale bar = 200 μm. (B) Quantification of GFP levels of *clec-60p::gfp*, *cmtr-1(jsn21);clec-60p::gfp*, *zip-2(tm4248);clec-60p::gfp*, and *zip-2(tm4248);cmtr-1(jsn21);clec-60p::gfp* worms. \*\*\**p*< 0.001 via t-test (*n* = 30 worms each). (C) Representative fluorescence images of N2, *cmtr-1(jsn21)*, *zip-2(tm4248)*, and *zip-2(tm4248);cmtr-1(jsn21)* worms incubated on *P. aeruginosa*-GFP for 24 hours at 25°C. Dotted outlines indicate worm positions. Scale bar = 200 μm. (D) Quantification of GFP levels of N2, *cmtr-1(jsn21)*, *zip-2(tm4248)*, and *zip-2(tm4248);cmtr-1(jsn21)* worms incubated on *P. aeruginosa*-GFP for 24 hours at 25°C. \*\*\**p*< 0.001 and \*\**p*< 0.01 via t-test (*n* = 30 worms each). (E) Representative survival plots of N2 and *cmtr-1(jsn21)* worms on *P. aeruginosa* PA14 at 25°C. \*\*\**p*< 0.001 for *cmtr-1(jsn21)* compared to N2 (*n* = 90 for N2 and 105 for *cmtr-1(jsn21)*). (F) Representative survival plots of *zip-2(tm4248)* and *zip-2(tm4248);cmtr-1(jsn21)* worms on *P. aeruginosa* PA14 at 25°C. \*\**p*< 0.01 for *zip-2(tm4248);cmtr-1(jsn21)* compared to *zip-2(tm4248)* (*n* = 90 per condition).

We next tested whether other major innate immune pathways were required for elevated immune activity in *cmtr-1(jsn21)* mutants. To this end, we crossed *cmtr-1(jsn21)* into mutants of the NSY-1/SEK-1/PMK-1 MAPK (Kim *et al*, 2002), KGB-1 (Kim *et al*, 2004), TGF-β/DBL-1 (Mallo *et al*, 2002), and TFEB/HLH-30 (Visvikis *et al*, 2014) pathways, and examined *clec-60* induction. Interestingly, *pmk-1(km25);cmtr-1(jsn21);clec-60p::gfp* worms exhibited severely reduced brood sizes, whereas *kgb-1(km21);cmtr-1(jsn21);clec-60p::gfp* worms were sterile, preventing meaningful evaluation of these mutants.

To circumvent this limitation, we examined the effects of *pmk-1* and *kgb-1* knockdown by RNAi. Although *cmtr-1(jsn21);clec-60p::gfp* worms developed on *pmk-1* and *kgb-1* RNAi, they exhibited either severely reduced brood sizes or complete sterility (Fig S5A). In contrast, knockdown of *pmk-1* or *kgb-1* did not affect the brood size of *clec-60p::gfp* control worms. Notably, *cmtr-1(jsn21)* mutants also displayed a modest reduction in brood size under control conditions, potentially reflecting a fitness trade-off associated with elevated immune activity, as reported previously (Foster *et al*, 2020). Despite the effects on brood size, knockdown of either *pmk-1* or *kgb-1* failed to suppress *clec-60p::gfp* induction in *cmtr-1(jsn21)* mutants (Fig S5B, S5C). Similarly, neither *dbl-1(nk3)* nor *hlh-30(tm1978)* suppressed *clec-60* induction in the *cmtr-1(jsn21)* background (Fig S6A, S6B). Collectively, these findings suggested that the enhanced immune response in *cmtr-1(jsn21)* mutants is unlikely to be mediated by the canonical immune pathways examined in this study.

### The GATA transcription factor ELT-2 mediates immune activation downstream of *cmtr-1(jsn21)*

GO analysis of upregulated genes revealed enrichment for transcription factors (Fig S3A), prompting us to identify transcriptional regulators driving immune activation in *cmtr-1(jsn21)* mutants. We conducted a feeding-based RNAi screen targeting 797 predicted transcription factors, representing 85% of factors listed in wTF3.0 (Fuxman Bass *et al*, 2016). Using *cmtr-1(jsn21);clec-60p::gfp* worms, we screened for knockdowns that lowered GFP expression and identified *elt-2* as the sole robust hit (Fig 5A).

**Figure 5:**
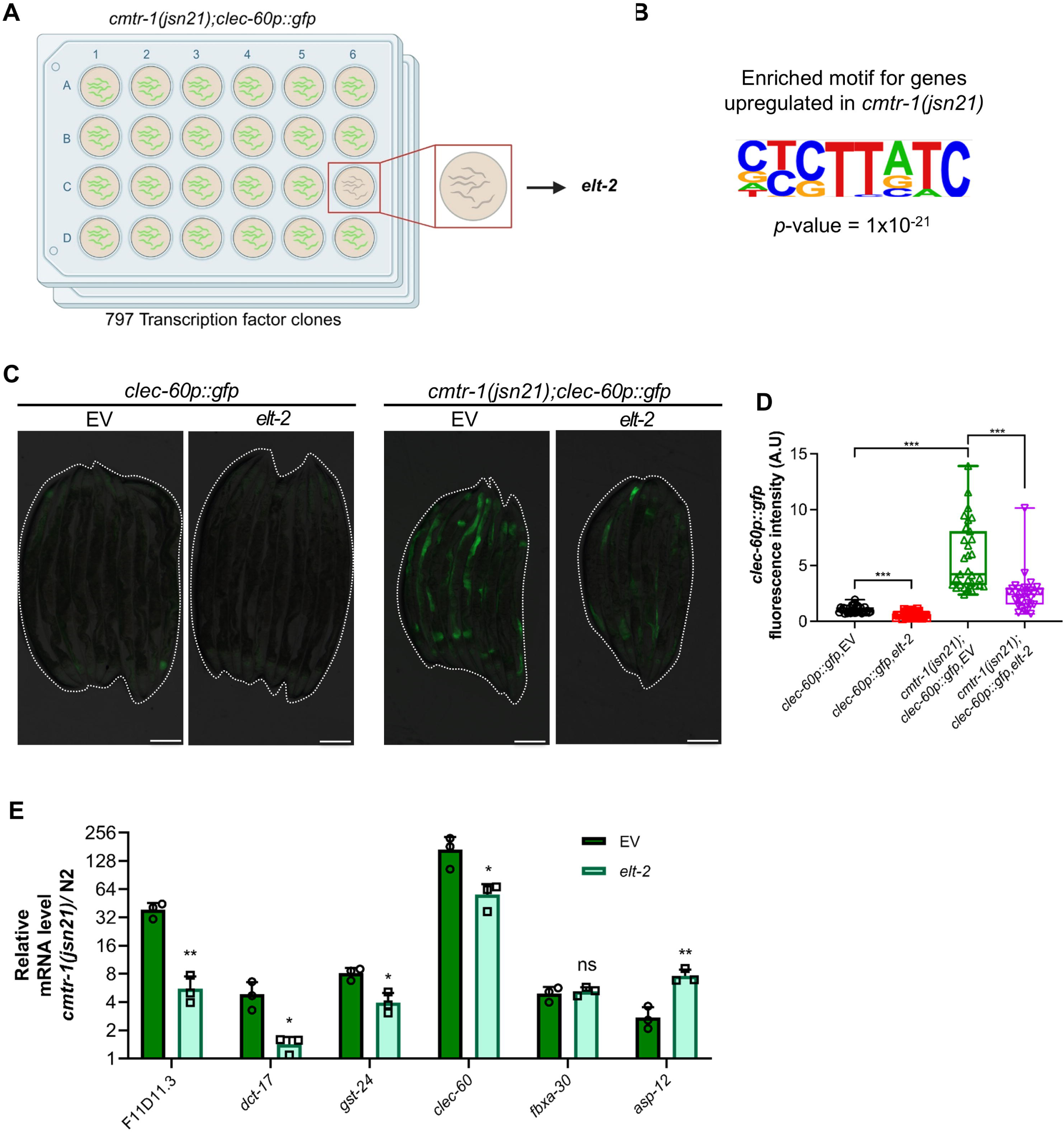
The GATA transcription factor ELT-2 mediates immune activation downstream of *cmtr-1(jsn21)* (A) Schematic representation of the feeding-based RNAi screen targeting 797 predicted transcription factors for suppressed GFP levels in *cmtr-1(jsn21);clec-60p::gfp* worms. The illustration was created using <u>BioRender</u>. (B) The most enriched binding motif in the promoters of 1217 genes upregulated in *cmtr-1(jsn21);clec-60p::gfp* worms. The *p-*value was derived from HOMER (a one-sided hypergeometric test). (C) Representative fluorescence images of *clec-60p::gfp* and *cmtr-1(jsn21);clec-60p::gfp* worms exposed to empty vector (EV) and *elt-2* RNAi. Dotted outlines indicate worm positions. Scale bar = 200 μm. (D) Quantification of GFP levels of *clec-60p::gfp* and *cmtr-1(jsn21);clec-60p::gfp* worms exposed to EV and *elt-2* RNAi. \*\*\**p*< 0.001 via t-test (*n* = 29-31 worms each). (E) qRT-PCR analysis of innate immune response genes in *cmtr-1(jsn21)* worms relative to N2 worms exposed to EV and *elt-2* RNAi. \*\**p*< 0.01, \**p*< 0.05, and ns, non-significant via t-test. Data represent the mean and standard deviation from three independent experiments.

To independently assess transcription factors that might regulate gene expression downstream of the *cmtr-1* mutation, we performed transcription factor motif enrichment analysis of the 1,217 upregulated genes using HOMER. Although several motifs were enriched, ELT-2 motifs showed the strongest enrichment (Fig 5B; Fig S7A). Consistently, *elt-2* RNAi markedly reduced GFP fluorescence in *cmtr-1(jsn21);clec-60p::gfp* worms (Fig 5C, 5D). To determine whether ELT-2 broadly regulates immune activation in *cmtr-1(jsn21)* mutants, we quantified mRNA levels of known ELT-2-dependent immune genes. The knockdown of *elt-2* reduced the expression of most of these genes in *cmtr-1(jsn21)* mutants (Fig 5E), indicating that ELT-2 mediated a substantial component of the immune activation induced by *cmtr-1* inhibition.

We next examined whether *cmtr-1* inhibition alters the nuclear abundance of ELT-2. Knockdown of *cmtr-1* did not affect the nuclear intensity of ELT-2::GFP (Fig S7B, S7C), suggesting that CMTR-1 inhibition likely modulates ELT-2 activity independently of changes in its nuclear localization or abundance.

### ELT-2 is required for enhanced pathogen resistance in *cmtr-1(jsn21)* mutants

We next investigated whether ELT-2 is required for the enhanced resistance of *cmtr-1(jsn21)* mutants to *P. aeruginosa* infection. The knockdown of *elt-2* increased gut colonization in both wild-type and *cmtr-1(jsn21)* worms (Fig 6A, 6B), demonstrating that ELT-2 contributes to pathogen resistance in each background. Consistent with its established role in immunity (Shapira *et al*, 2006), *elt-2* RNAi reduced the survival of wild-type worms during infection (Fig 6C). Importantly, *elt-2* knockdown abolished the enhanced survival phenotype of *cmtr-1(jsn21)* mutants, such that *cmtr-1(jsn21)* worms on *elt-2* RNAi no longer survived better than wild-type worms under the same conditions (Fig 6C). Collectively, these results demonstrated that the enhanced immune response and pathogen resistance of *cmtr-1(jsn21)* mutants are mediated by ELT-2.

**Figure 6:**
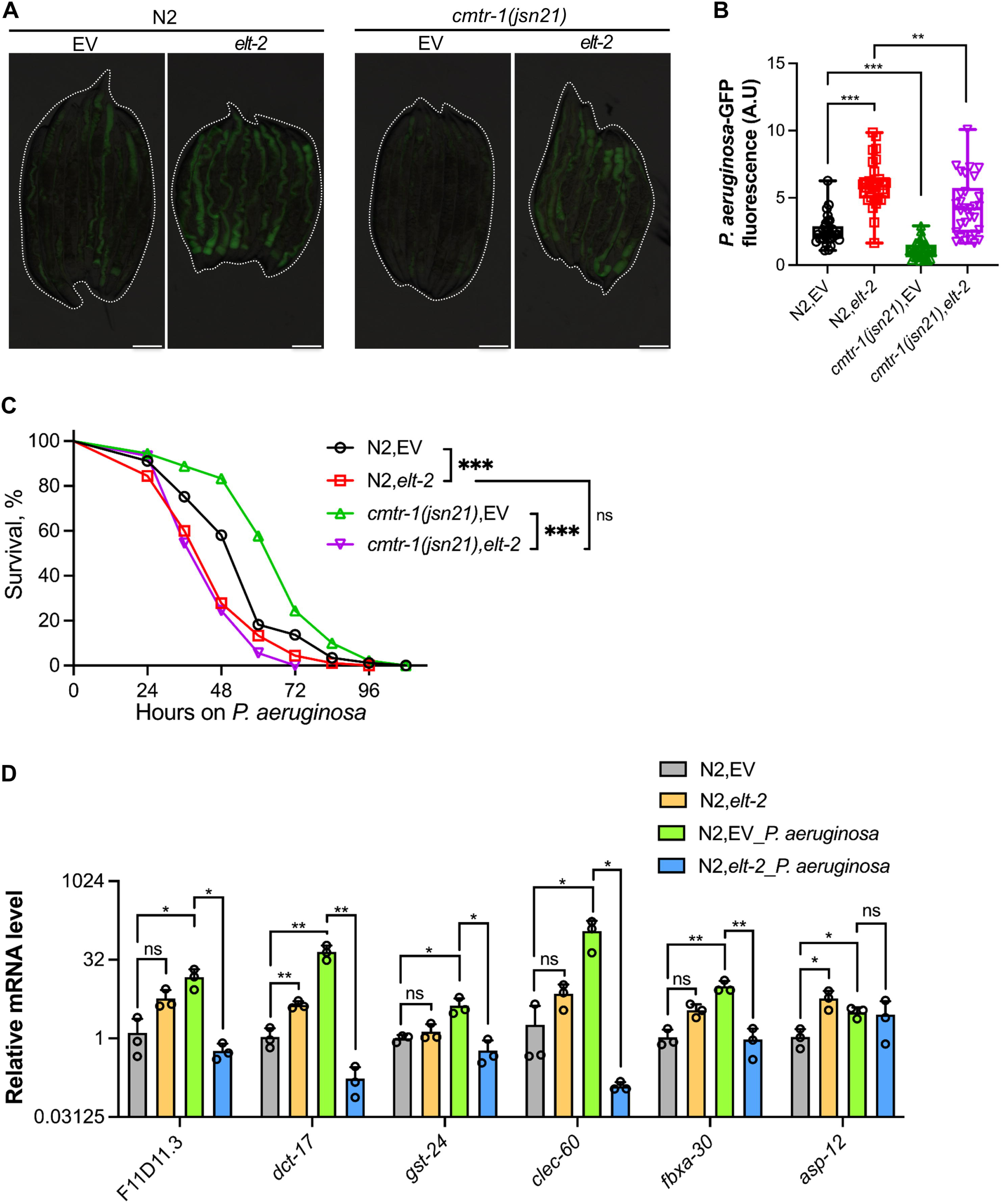
ELT-2 is required for enhanced pathogen resistance in *cmtr-1(jsn21)* mutants. (A) Representative fluorescence images of N2 and *cmtr-1(jsn21)* worms on *P. aeruginosa*-GFP for 24 hours at 25°C after treatment with empty vector (EV) and *elt-2* RNAi. Dotted outlines indicate worm positions. Scale bar = 200 μm. (B) Quantification of GFP levels of N2 and *cmtr-1(jsn21)* worms on *P. aeruginosa*-GFP for 24 hours at 25°C after treatment with EV and *elt-2* RNAi. \*\*\**p*< 0.001 and \*\**p*< 0.01 (*n* = 30 worms each). (C) Representative survival plots of N2 and *cmtr-1(jsn21)* worms on *P. aeruginosa* PA14 at 25°C after treatment with EV and *elt-2* RNAi. \*\*\**p*< 0.001 for both N2 and *cmtr-1(jsn21)* on *elt-2* RNAi as compared to N2 and *cmtr-1(jsn21)* on EV, respectively. *p* is non-significant between *cmtr-1(jsn21)* on *elt-2* RNAi and N2 on *elt-2* RNAi (*n* = 90 for N2 on EV and *elt-2* RNAi, 92 for *cmtr-1(jsn21)* on EV, and 60 for *cmtr-1(jsn21)* on *elt-2* RNAi). (D) qRT-PCR analysis of innate immune response genes in N2 worms exposed to EV and *elt-2* RNAi with and without *P. aeruginosa* PA14 infection for 24 hours at 25°C. \*\**p*< 0.01, \**p*< 0.05, and ns, non-significant via t-test. Data represent the mean and standard deviation from three independent experiments.

We next asked whether the immune genes upregulated in *cmtr-1(jsn21)* mutants are also induced during *P. aeruginosa* infection. Notably, all tested genes that were upregulated in *cmtr-1(jsn21)* mutants were also induced following *P. aeruginosa* infection (Fig 6D), suggesting that *cmtr-1* inhibition primes components of the immune response normally activated during pathogen infection. Importantly, induction of these genes during *P. aeruginosa* infection was fully suppressed by *elt-2* knockdown (Fig 6D), indicating that ELT-2 regulates these immune genes in response to both pathogen infection and *cmtr-1* inhibition. In contrast, *elt-2* knockdown did not reduce the basal expression of these genes prior to pathogen exposure (Fig 6D), suggesting that ELT-2 is dispensable for their constitutive expression under non-infectious conditions. Collectively, these findings demonstrated that ELT-2 controls a common immune transcriptional program activated both during *P. aeruginosa* infection and upon inhibition of CMTR-1.

## Discussion

Surveillance of core cellular processes is a critical component of host defense against bacterial infection (Willmann & Moita, 2025). Because diverse pathogens disrupt distinct aspects of cellular homeostasis, including protein translation, mitochondrial function, and proteasome activity, hosts have evolved multiple mechanisms to sense these perturbations and initiate protective immune responses. For example, inhibition of host translation triggers immunity through the bZIP transcription factor ZIP-2 and the CCAAT/enhancer-binding protein CEBP-2 (Dunbar *et al*, 2012; Ghosh & Singh, 2024; Reddy *et al*, 2016), while mitochondrial and proteasomal stress activate immune responses via ATFS-1 and SKN-1, respectively (Kahn *et al*, 2007; Papp *et al*, 2012; Pellegrino *et al*, 2014).

In this study, we identify inhibition of the conserved mRNA cap methyltransferase CMTR-1 as a previously unrecognized trigger of innate immune activation, acting through the GATA transcription factor ELT-2. Notably, ELT-2 was recently shown to mediate lysosomal surveillance (Li *et al*, 2025), raising the possibility that multiple damage-sensing pathways converge on this transcription factor to coordinate immune activation. Interestingly, in a recent study, knockout of *cmtr-1* was shown to induce immune responses associated with intracellular pathogen defense (Clemens *et al*, 2026). In agreement with these findings, we also observed upregulation of several *pals* genes (Table S1), which are established components of the intracellular pathogen response pathway (Reddy *et al*, 2017).

In higher eukaryotes, the ribose 2’-O-methylation of the first transcribed nucleotide, catalyzed by CMTR1 (Bélanger *et al*, 2010), acts in concert with the 5’-methylated guanosine cap to distinguish self (methylated) from non-self (unmethylated) RNA (Schuberth-Wagner *et al*, 2015). CMTR1 is upregulated in response to several viral infections and promotes the translation of interferon-stimulated genes, underscoring its relevance to antiviral immunity (Lukoszek *et al*, 2024; Williams *et al*, 2020). However, its role in bacterial infection has not been explored. Here, we demonstrate that loss of *cmtr-1* in *C. elegans* activates ELT-2-dependent immune responses, which enhance resistance to the bacterial pathogen *P. aeruginosa*. Whether bacterial infection directly modulates CMTR-1 activity will require further investigation.

Transcriptomic analysis revealed that *cmtr-1* loss-of-function reduces the expression of numerous ribosomal protein genes, suggesting impaired translation. This observation is consistent with previous studies in murine embryonic stem cells, which show that CMTR1 binds to the transcription start sites of ribosomal protein genes, thereby regulating their expression and overall translational capacity (Liang *et al*, 2022). Because translation inhibition is known to activate immunity via ZIP-2 (Dunbar *et al*, 2012; Ghosh & Singh, 2024), we tested whether this pathway contributed to the immune response in *cmtr-1(jsn21)* mutants. Our results show that ZIP-2 is not required for immune activation following CMTR-1 inhibition. Instead, immune induction in the mutant is mediated by ELT-2. However, the molecular mechanism by which CMTR-1 loss activates the ELT-2 pathway remains unresolved. Future studies will be needed to define how perturbation of mRNA cap methylation is sensed and coupled to ELT-2-dependent immune activation.

## Materials and methods

### Bacterial strains

The following bacterial strains were used in the current study: *Escherichia coli* OP50, *E. coli* HT115(DE3), *Pseudomonas aeruginosa* PA14, and *P. aeruginosa* PA14 expressing green fluorescent protein (*P. aeruginosa* PA14-GFP). GFP expression in *P. aeruginosa* PA14-GFP originates from plasmid pRR54GFP19-1, as described previously (Tan *et al*, 1999). Cultures of *E. coli* OP50, *E. coli* HT115(DE3), and *P. aeruginosa* PA14 were grown in Luria-Bertani (LB) broth at 37°C, while cultures of *P. aeruginosa* PA14-GFP were grown in LB broth with 50 µg/mL kanamycin at 37°C.

### *C. elegans* strains and growth conditions

*C. elegans* hermaphrodites were maintained at 20°C on nematode growth medium (NGM) plates seeded with *E. coli* OP50 as the food source unless otherwise indicated. Bristol N2 hermaphrodites were used as the wild-type control unless otherwise indicated. The following strains were used in the study: JIN810 *clec-60p::gfp*, JIN1375 *hlh-30(tm1978)*, KU25 *pmk-1(km25)*, NU3 *dbl-1(nk3)*, KU21 *kgb-1(km21)*, *zip-2(tm4248)*, and SD1949 *glo-4(ok623);gaIs290 [elt-2::TY1::EGFP::3xFLAG(92C12)+unc-119(+)].* Some of the strains were obtained from the Caenorhabditis Genetics Center (University of Minnesota, Minneapolis, MN), while *zip-2(tm4248*) was obtained from the National BioResource Project (NBRP), Japan. The following strains were generated in this study: JSJ21 *cmtr-1(jsn21);clec-60p::gfp*, *cmtr-1(jsn21);clec-60p::gfp;jsnEx10[sur-5p::cmtr-1 + myo-3p::mCherry]*, and *cmtr-1(jsn21);clec-60p::gfp;jsnEx11[vha-6p::cmtr-1 + myo-3p::mCherry].* The following strains were obtained by standard genetic crosses: *pmk-1(km25);clec-60p::gfp, kgb-1(km21);clec-60p::gfp, dbl-1(nk3);clec-60p::gfp, hlh-30(tm1978);clec-60p::gfp, zip-2(tm4248);clec-60p::gfp*, *pmk-1(km25);cmtr-1(jsn21);clec-60p::gfp*, *kgb(km21);cmtr-1(jsn21);clec-60p::gfp, dbl-1(nk3);cmtr-1(jsn21);clec-60p::gfp; hlh-30(tm1978);cmtr-1(jsn21);clec-60p::gfp, zip-2(tm4248);cmtr-1(jsn21);clec-60p::gfp*. For performing *P. aeruginosa* PA14-GFP colonization assays, *cmtr-1(jsn21)*, *cmtr-1(jsn21);jsnEx10[sur-5p::cmtr-1 + myo-3p::mCherry]*, *cmtr-1(jsn21);jsnEx11[vha-6p::cmtr-1 + myo-3p::mCherry]*, and *zip-2(tm4248);cmtr-1(jsn21)* strains were generated by removing the *clec-60p::gfp* transgene through crosses with N2 males. Genotyping primer sequences are provided in Table S2.

### RNA interference (RNAi)

RNAi was performed by feeding worms with *E. coli* HT115(DE3) expressing double-stranded RNA homologous to a target *C. elegans* gene, as described earlier (Ghosh & Singh, 2024, 2025). Briefly, *E. coli* with the relevant vectors were grown overnight in LB broth containing ampicillin (100 μg/mL) at 37°C and plated onto RNAi NGM plates containing 100 μg/mL ampicillin and 3 mM isopropyl β-D-thiogalactoside (IPTG). The RNAi-expressing bacteria were allowed to grow overnight at 37°C on RNAi plates. Worms were synchronized on RNAi plates, and eggs were allowed to develop at 20°C for 96 hours. RNAi clones were obtained from the Ahringer RNAi library and verified by sequencing.

### Transcription factor library screening

*E. coli* HT115(DE3) carrying RNAi clones for 797 predicted transcription factors were grown overnight at 37°C in LB broth containing ampicillin (100 μg/ml) in 96-well plates. The list of transcription factors was obtained from an earlier study (Fuxman Bass *et al*, 2016), and the transcription factor sub-library was created from the Ahringer RNAi library. A 30 μL aliquot of each culture was seeded onto 24-well RNAi NGM plates containing 100 μg/mL ampicillin and 3 mM IPTG. RNAi-expressing bacteria were grown overnight at 37°C. For screening, *cmtr-1(jsn21);clec-60p::gfp* worms were synchronized by bleaching in an alkaline bleach solution and then incubated in M9 buffer for 22 hours at room temperature to obtain synchronized L1 larvae. Approximately 25-30 synchronized L1 larvae were transferred to each RNAi well and incubated at 20°C. After 96 hours, wells were examined for transcription factors whose knockdown reduced *clec-60p::gfp* expression. Knockdown of certain transcription factors, including *elt-2*, caused developmental arrest. For these clones, *cmtr-1(jsn21);clec-60p::gfp* worms were grown on empty-vector bacteria until the L4 stage and then transferred to the respective transcription factor RNAi plates. GFP fluorescence was monitored after 48 hours of transferring the L4s.

### Binding motif enrichment analysis

Binding motif analysis for genes upregulated in *cmtr-1(jsn21)* worms was performed using HOMER (Heinz *et al*, 2010). The 1217 genes upregulated in *cmtr-1(jsn21)* worms were used as the input dataset in HOMER. The *C. elegans* genes and promoters databases were downloaded using Perl, and the enrichment analysis was performed using the findMotifs.pl script as described previously (Li *et al*, 2025).

### Forward genetic screens for mutants with enhanced *clec-60p::gfp* expression levels

Ethyl methanesulfonate (EMS) mutagenesis screens (Singh, 2021) were performed using the *clec-60p::gfp* strain. Approximately 2,500 synchronized late L4 larvae were treated with 50 mM EMS for 4 hours and washed three times with M9 medium. EMS-treated worms (P0 generation) were transferred to 9-cm NGM plates seeded with *E. coli* OP50 and allowed to lay eggs (F1 progeny) overnight. P0 worms were then removed by washing with M9 medium, leaving F1 eggs attached to the bacterial lawn. F1 eggs were allowed to reach adulthood. Adult F1 worms were bleached to obtain F2 eggs, which were transferred to *E. coli* OP50 plates and incubated at 20°C for 96 hours. Plates were then screened for worms with enhanced GFP expression. Approximately 50,000 haploid genomes were screened, and one fertile mutant was isolated. The mutant was backcrossed six times with the parental *clec-60p::gfp* strain before analysis.

### Whole-genome sequencing (WGS) and data analysis

Genomic DNA was isolated as described previously (Gokul & Singh, 2022; Ravi *et al*, 2023). Briefly, mutant worms were grown at 20°C on NGM plates seeded with *E. coli* OP50 until starvation. Four 9-cm plates were used for each strain to obtain a sufficient number of worms. Worms were rinsed from the plates with M9, washed three times, and then incubated in M9 with rotation for 2 hours to remove intestinal bacteria. Worms were washed three additional times with distilled water and stored at −80°C until genomic DNA extraction. Genomic DNA was extracted using the Gentra Puregene Kit (Qiagen, Netherlands). DNA libraries were prepared using standard Illumina protocols, and sequencing was performed on an Illumina NovaSeq 6000 platform using 150 bp paired-end reads at the National Genomics Core, National Institute of Biomedical Genomics, Kalyani, India.

WGS data were analyzed using the Galaxy web platform, as described earlier (Ghosh & Singh, 2025; Gokul & Singh, 2022). Briefly, forward and reverse FASTQ reads, the *C. elegans* reference genome FASTA file (ce11M.fa), and the gene annotation file (SnpEff4.3 WBcel235.86) were input into the Galaxy workflow. Low-quality read ends were trimmed using Sickle, and trimmed reads were aligned to the reference genome with BWA-MEM. Duplicate reads were filtered using MarkDuplicates. Variants, including single-nucleotide polymorphisms, small insertions and deletions, multi-nucleotide polymorphisms, and complex events smaller than typical short-read alignment length, were identified with FreeBayes. To subtract background variation, WGS data from mutants isolated in an independent *clec-60p::gfp* screen (Ghosh & Singh, 2025) were used. The SnpEff4.3 WBcel235.86 gene annotation file was used to annotate and predict the effects of genetic variants (such as amino acid changes). Linkage maps for the mutant were generated using the resulting variants.

### *C. elegans* slow-killing assays on *P. aeruginosa* PA14

Full-lawn slow-killing assays of *C. elegans* exposed to *P. aeruginosa* PA14 were performed as described previously (Das *et al*, 2024; Ghosh & Singh, 2024). Briefly, bacterial cultures were prepared by inoculating individual bacterial colonies of *P. aeruginosa* into 3 mL of LB broth and growing for 10-12 hours at 37°C with shaking. Bacterial lawns were prepared by spreading 20 µL of the culture onto the entire surface of 3.5-cm-diameter modified NGM agar plates (0.35% instead of 0.25% peptone). Plates were incubated at 37°C for 10-12 hours and then cooled to room temperature for at least 30 minutes before adding synchronized 1-day-old adult worms. Killing assays were conducted at 25°C, and live worms were transferred daily to fresh *P. aeruginosa* plates. Worms were scored at the indicated times and considered dead when unresponsive to touch. Killing assays upon the knockdown of *eif-2α* and *ifg-1* were carried out as described previously (Ghosh & Singh, 2024). At least three independent biological replicates were performed for each condition. Detailed statistical analyses for all survival assays are provided in Table S3.

### *C. elegans* fast-killing assays on *P. aeruginosa* PA14

Fast-killing assays using *P. aeruginosa* PA14 were performed as described previously (Tan *et al*, 1999), with slight modifications. The concentration of glucose and sorbitol were 20% and 1.5 M, respectively, in the fast-killing assay plates. *P. aeruginosa* PA14 was grown in 2 mL of LB broth at 37°C for 10-12 hours with shaking. Subsequently, 5 µL of the culture was spread onto peptone-glucose-sorbitol (PGS) agar plates. Following inoculation, the plates were incubated at 37°C for 24 hours and then maintained at room temperature for an additional 12 hours before use. Thirty synchronized L4-stage worms from each strain were transferred onto the assay plates, which were then maintained at 25°C. Worm survival was monitored hourly for the first 4 hours following transfer, with a final assessment performed at 24 hours. Three independent biological replicates were conducted for each strain. Detailed statistical analyses for all survival assays are provided in Table S3.

### *P. aeruginosa*-GFP colonization assay

*P. aeruginosa* PA14-GFP colonization assays were performed as described earlier (Ghosh & Singh, 2024; Rao *et al*, 2025). Briefly, bacterial cultures were prepared by inoculating individual bacterial colonies of *P. aeruginosa* PA14-GFP into 3 mL LB broth containing 50 μg/mL kanamycin and growing for 10-12 hours at 37°C with shaking. Bacterial lawns were prepared by spreading 20 µL of culture onto the entire surface of 3.5-cm-diameter modified NGM agar plates (0.35% instead of 0.25% peptone) containing 50 μg/mL kanamycin. Plates were incubated at 37°C for 12 hours and then cooled to room temperature for at least 30 minutes before seeding with 1-day-old adult gravid adults. Assays were performed at 25°C. At the indicated times, worms were picked under a non-fluorescent stereomicroscope and imaged within 5 minutes on a fluorescence microscope. At least three biological replicates were performed for each condition.

### Bacterial killing

Bacterial killing was performed as described previously (Gahlot *et al*, 2025). Briefly, *E. coli* OP50 was cultured in 50 mL LB broth at 37°C for 24 hours. Cultures were centrifuged at 3,000 × *g* for 30 minutes, the supernatant was removed, and the pellets were resuspended in 100 mL of autoclaved water containing 250 μg/mL kanamycin. The suspension was split equally into two 250-mL flasks and incubated with shaking at 37°C for an additional 24 hours. To confirm bacterial killing, 1 mL of culture was washed three times with sterile water and plated on LB agar. Plates were incubated at 37°C for 24 hours and examined for viable colonies. In the absence of colonies, bacteria were deemed effectively killed and stored at 4°C. Prior to use, killed bacteria were washed three times with 100 mL sterile water and concentrated 20-fold before being seeded onto NGM plates with or without 50 μg/mL 5-fluorodeoxyuridine (FUdR).

### *C. elegans* longevity assays in the presence of FUdR

Lifespan assays were performed as described previously (Das *et al*, 2024, 2025). Assays were conducted on both live and kanamycin-killed *E. coli* OP50 in the presence of 50 µg/mL FUdR. *E. coli* OP50 was killed as described above. Worms were synchronized on plates containing either live or dead *E. coli* without FUdR and incubated at 20°C. At the late L4 larval stage, worms were transferred to FUdR-containing plates seeded with either live or dead OP50, respectively, and incubated at 20°C. Worms were scored every other day as live or dead. Worms that failed to display touch-provoked movement were scored as dead. Worms that crawled off plates were censored. Young adults were designated as day 0 for lifespan analysis. Three independent biological replicates were performed. Detailed statistical analyses for all survival assays are provided in Table S3.

### *C. elegans* longevity assays in the absence of FUdR

For lifespan assays performed without FUdR, 90-100 synchronized late L4-stage worms raised on either live or kanamycin-killed *E. coli* OP50 were transferred to fresh NGM plates seeded with the corresponding bacterial condition. The plates were maintained at 20°C throughout the assay. During the reproductive period, live worms were transferred daily to fresh *E. coli* OP50 plates for the first 6-7 days until egg laying ceased. Worms that failed to exhibit touch-provoked movement were scored as dead, whereas worms that crawled off the plates were censored from the analysis. Young adult worms were designated as day 0 for lifespan analysis. Two independent biological replicates were performed. Detailed statistical analyses for all survival assays are provided in Table S3.

### *C. elegans* RNA isolation and quantitative reverse transcription-PCR

Wild-type *clec-60p::gfp* and *cmtr-1(jsn21);clec-60p::gfp* worms were synchronized by egg-laying. Approximately 50 gravid adult worms for *clec-60p::gfp* and 100 for the *cmtr-1(jsn21);clec-60p::gfp* were transferred to *E. coli* OP50 plates and allowed to lay eggs for 4 hours. Gravid adults were removed, and eggs were allowed to develop at 20°C for 96 hours. For experiments involving *elt-2* RNAi, both N2 and *cmtr-1(jsn21)* worms were synchronized on empty vector (EV) control bacteria until the L4 stage (48 hours after egg laying for N2; 54 hours for *cmtr-1(jsn21)*). Synchronized L4 larvae were collected in M9, washed twice, and transferred to 9-cm *elt-2* RNAi plates and grown to 1-day-old adults. Control worms were grown on EV RNAi plates till they grew to 1-day-old adults. For experiments involving pathogen exposure following RNAi treatment, EV- and *elt-2* RNAi-treated N2 worms were transferred to 9-cm slow-killing assay plates seeded with 120 µL of an overnight culture of *P. aeruginosa* PA14. The plates were then incubated at 25°C for 24 hours. Subsequently, worms were collected, washed in M9, and frozen in TRIzol reagent (Life Technologies, Carlsbad, CA, USA). Total RNA was extracted using the RNeasy Plus Universal Kit (Qiagen, the Netherlands). One microgram of total RNA was reverse-transcribed with random primers using the PrimeScript 1st Strand cDNA Synthesis Kit (TaKaRa). qRT-PCR was performed using TB Green fluorescence (TaKaRa) on a MasterCycler EP Realplex 4 (Eppendorf) in 96-well format. Fifteen-microliter reactions were set up according to the manufacturer’s instructions. Relative transcript levels were calculated using the comparative *Ct*(2^-ΔΔ^*Ct*) method and normalized to pan-actin (*act-1,3,4*) as described previously (Ghosh & Singh, 2024). All samples were analyzed in technical triplicate and repeated in at least three independent biological replicates. Primer sequences are provided in Table S2.

### RNA sequencing and data analysis

For RNA sequencing, worm synchronization and RNA extraction for *clec-60p::gfp* and *cmtr-1(jsn21);clec-60p::gfp* worms were performed as described above for qRT-PCR. Three biological replicates were prepared for each condition. Library preparation and sequencing were performed at the National Genomics Core, National Institute of Biomedical Genomics, Kalyani, India, using an Illumina NovaSeq 6000 platform with 150 bp paired-end reads.

RNA sequencing data were analyzed on the Galaxy web platform (https://usegalaxy.org/) following established procedures (Ghosh & Singh, 2024). Briefly, paired-end reads were quality trimmed with Trimmomatic, aligned to the *C. elegans* reference genome (WS220) using STAR, and quantified using htseq-count. Differential gene expression analysis was performed using DESeq2. Genes with ≥2-fold differential expression and P < 0.01 were considered significantly altered. Gene Ontology enrichment was assessed using the DAVID Bioinformatics Database (https://david.ncifcrf.gov/tools.jsp).

### Plasmid constructs and generation of transgenic *C. elegans*

For pan-tissue rescue of *cmtr-1(jsn21)*, the wild-type copy of *cmtr-1* was expressed under the *sur-5* promoter. The *sur-5* promoter was used instead of the native *cmtr-1* promoter because the latter was found to be difficult to clone. The 1,005 bp upstream region of *sur-5* was amplified from N2 genomic DNA and cloned into the pPD95_77 plasmid using HindIII and SalI. The 2,779 bp *cmtr-1* cDNA was amplified from N2 cDNA and cloned into the *sur-5p* construct using XbaI and KpnI. The resulting *sur-5p::cmtr-1* plasmid was microinjected into *cmtr-1(jsn21);clec-60p::gfp* worms at 50 ng/μL along with pCFJ104 (*myo-3p::mCherry*) at 25 ng/μL as a co-injection marker. cDNA was synthesized using the PrimeScript 1st Strand cDNA Synthesis Kit (TaKaRa) with oligo-dT primers.

For intestine-specific rescue, *cmtr-1* cDNA was cloned into pDB24 (*vha-6p::mCherry*) using XbaI and KpnI. The pDB24, which contains the intestine-specific promoter *vha-6* in the pPD95_77 backbone, has been described earlier (Ghosh & Singh, 2025). The resulting *vha-6p::cmtr-1* construct was co-injected into *cmtr-1(jsn21);clec-60p::gfp* worms at 50 ng/μL with pCFJ104 (*myo-3p::mCherry*) at 25 ng/μL. Plasmids were maintained as extrachromosomal arrays *jsnEx10[sur-5p::cmtr-1 + myo-3p::mCherry]* and *jsnEx11[vha-6p::cmtr-1 + myo-3p::mCherry]*. At least two independent transgenic lines were generated and examined for each construct. Images were obtained from one of the stable lines. Primer sequences are provided in Table S2.

### Brood size assay

Brood size assays were performed using *clec-60p::gfp* and *cmtr-1(jsn21);clec-60p::gfp* strains. Worms were synchronized on EV control, *pmk-1*, or *kgb-1* RNAi plates until the late L4 larval stage. Subsequently, 10 individual worms from each strain and condition were transferred onto separate RNAi plates and allowed to lay eggs for 24 hours. Each worm was transferred daily to freshly prepared RNAi plates until egg laying ceased. Progeny on each plate were allowed to develop until the L3 larval stage, at which point they were counted. The total brood size for each worm was calculated by summing the progeny produced across all plates for the respective strain and condition.

### Fluorescence imaging of *C. elegans*

Fluorescence imaging was performed as described previously (Das *et al*, 2025; Ghosh & Singh, 2024). Briefly, worms were picked under a non-fluorescent stereomicroscope to prevent selection bias. Worms were anesthetized in M9 buffer containing 50 mM sodium azide and mounted on 2% agarose pads. Fluorescence imaging was conducted using either a Nikon SMZ-1000 or SMZ18 stereomicroscope. Fluorescence intensity was quantified using ImageJ.

For ELT-2::GFP imaging, fluorescence images were acquired using a Zeiss confocal microscope (LSM 710). Nuclear fluorescence intensity was manually quantified using ImageJ in both anterior and posterior intestinal nuclei from 10-12 individual worms for each condition.

### Quantification and statistical analysis

All statistical analyses were performed with Prism 8 (GraphPad), as described previously (Ghosh & Singh, 2025). Data are presented as mean ± standard deviation. Statistical significance was assessed using an unpaired, two-tailed *t*-test, with *p* < 0.05 considered significant. In all figures, asterisks indicate significance levels as follows: \**p* < 0.05; \*\**p* < 0.01; and \*\*\**p* < 0.001, relative to the corresponding controls. Survival analyses were conducted using the Kaplan-Meier method, with statistical differences assessed using the log-rank test. All experiments were performed at least three times independently unless otherwise indicated.

## Acknowledgments

We thank the Caenorhabditis Genetics Center [funded by the NIH Office of Research Infrastructure Programs (P40 OD010440)] for providing the strains used in this study. We thank Ravi for cloning the *sur-5* promoter and Kaustubh Prakash for assistance with the HOMER installation and binding motif analysis. We also thank Dr. Mahak Sharma’s laboratory (IISER Mohali) for access to the confocal microscopy facility.

## Funding

This work was supported by the following grants: Har-Gobind Khorana-Innovative Young Biotechnologist Fellowship to J.S. (File No. HRD-17011/2/2023-HRD-DBT) and Ramalingaswami Re-entry Fellowship to J.S. (Ref. No. BT/RLF/Re-entry/50/2020) awarded by the Department of Biotechnology, India; STARS grant to J.S. (File No. MoE-STARS/STARS-2/2023-0116) awarded by the Ministry of Education, India; Anusandhan National Research Foundation (ANRF) Core Research Grant to J.S. (Ref. No. CRG/2023/001136) awarded by DST, India; Research Grant to J.S. (Ref. No. 37/1741/23/EMR-II) awarded by the Council of Scientific & Industrial Research (CSIR), India; and Indian Institute of Science Education and Research Mohali intramural funds. The funders had no role in study design, data collection and analysis, decision to publish, or preparation of the manuscript.

## Author Contributions

A.G. and J.S. conceived and designed the experiments. A.G. performed the experiments. A.G. and J.S. analyzed the data and wrote the paper.

## Competing interests

The authors have declared that no competing interests exist.

## Data availability

Whole-genome sequencing data for the JSJ21 strain have been deposited in the Sequence Read Archive under BioProject ID PRJNA1376507. RNA-sequencing data for *clec-60p::gfp* and *cmtr-1(jsn21);clec-60p::gfp* worms are available in the Sequence Read Archive under BioProject ID PRJNA1376512. All other data generated or analyzed in this study are provided within the manuscript and its supplementary files.

## Supplementary tables

**Table S1:** Upregulated and downregulated genes in *cmtr-1(jsn21);clec-60p::gfp* worms versus *clec-60p::gfp* worms. Genes exhibiting at least two-fold change and *p*-value <0.01 were considered differentially expressed. A separate Excel file is provided.

**Table S2:** Primers used in the study. A separate Excel file is provided.

**Table S3:** Statistical analysis of survival curves from three independent experiments for each condition. A separate Excel file is provided.

## Notes

### Competing Interest Statement

The authors have declared no competing interest.

### Summary of Updates

Data were added to clarify the role of translation inhibition in immune responses following CMTR-1 inhibition, the effect of FUdR on the lifespan of cmtr-1 mutants, and the effects of pmk-1 and kgb-1 knockdown on clec-60 expression in cmtr-1 mutants. Supplemental files updated.

