## Supporting Information for "RNA methyltransferase CMTR-1 inhibition activates a GATA transcription factor-mediated protective immune response"

**Supplementary Figures**

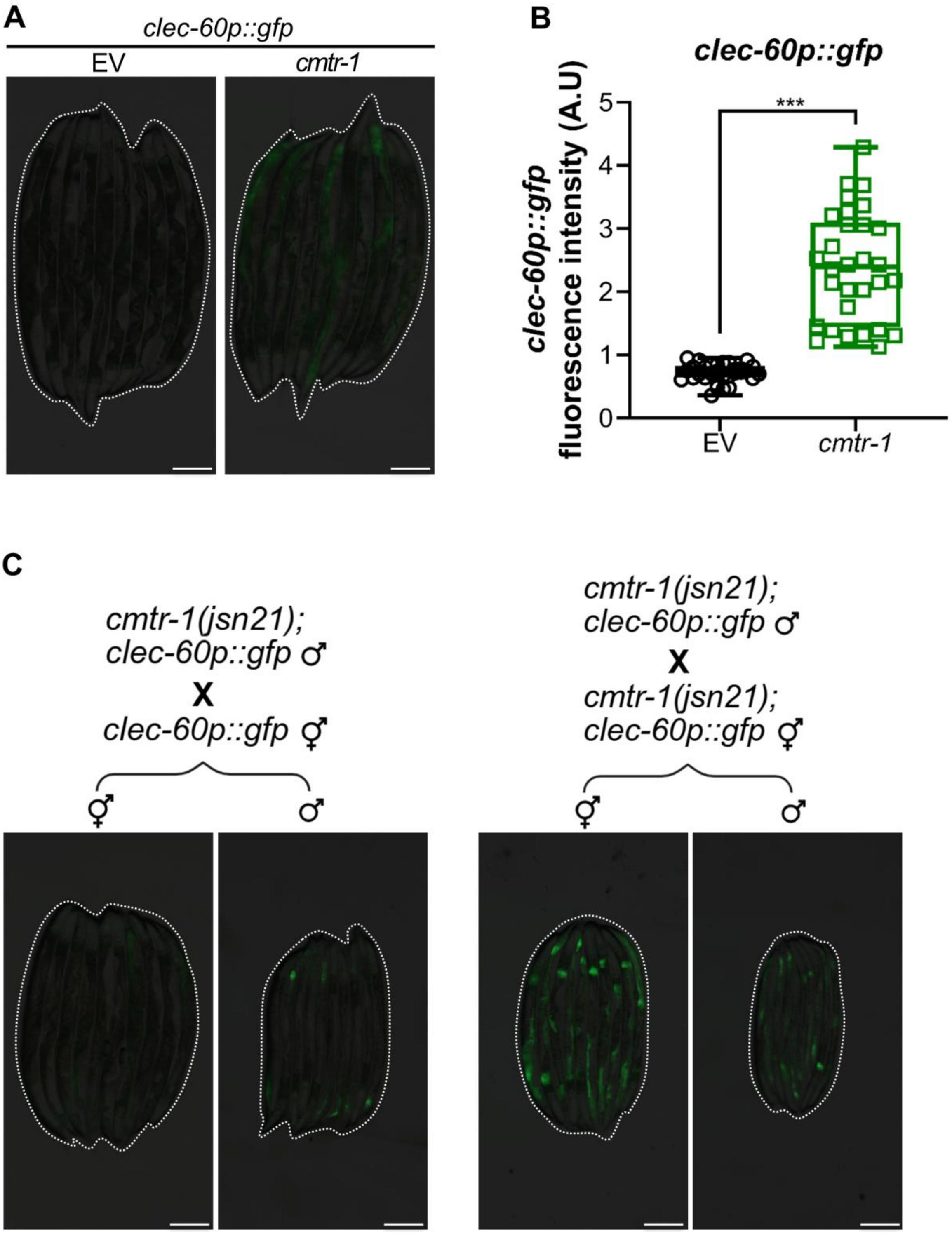


**Figure S1: Loss of *cmtr-1* enhances *clec-60p::gfp* levels**

(A) Representative fluorescence images of *clec-60p::gfp* worms exposed to empty vector (EV) and *cmtr-1* RNAi. Dotted outlines indicate worm positions. Scale bar = 200 µm.

(B) Quantification of GFP levels of *clec-60p::gfp* worms exposed to EV and *cmtr-1* RNAi. ***$p < 0.001$ via t-test ($n = 30$ worms each).

(C) Representative fluorescence images of the male and hermaphrodite F1 progeny of the shown genetic crosses. Dotted outlines indicate worm positions. The illustration of the genetic crosses was created using BioRender. Scale bar = 200 µm.

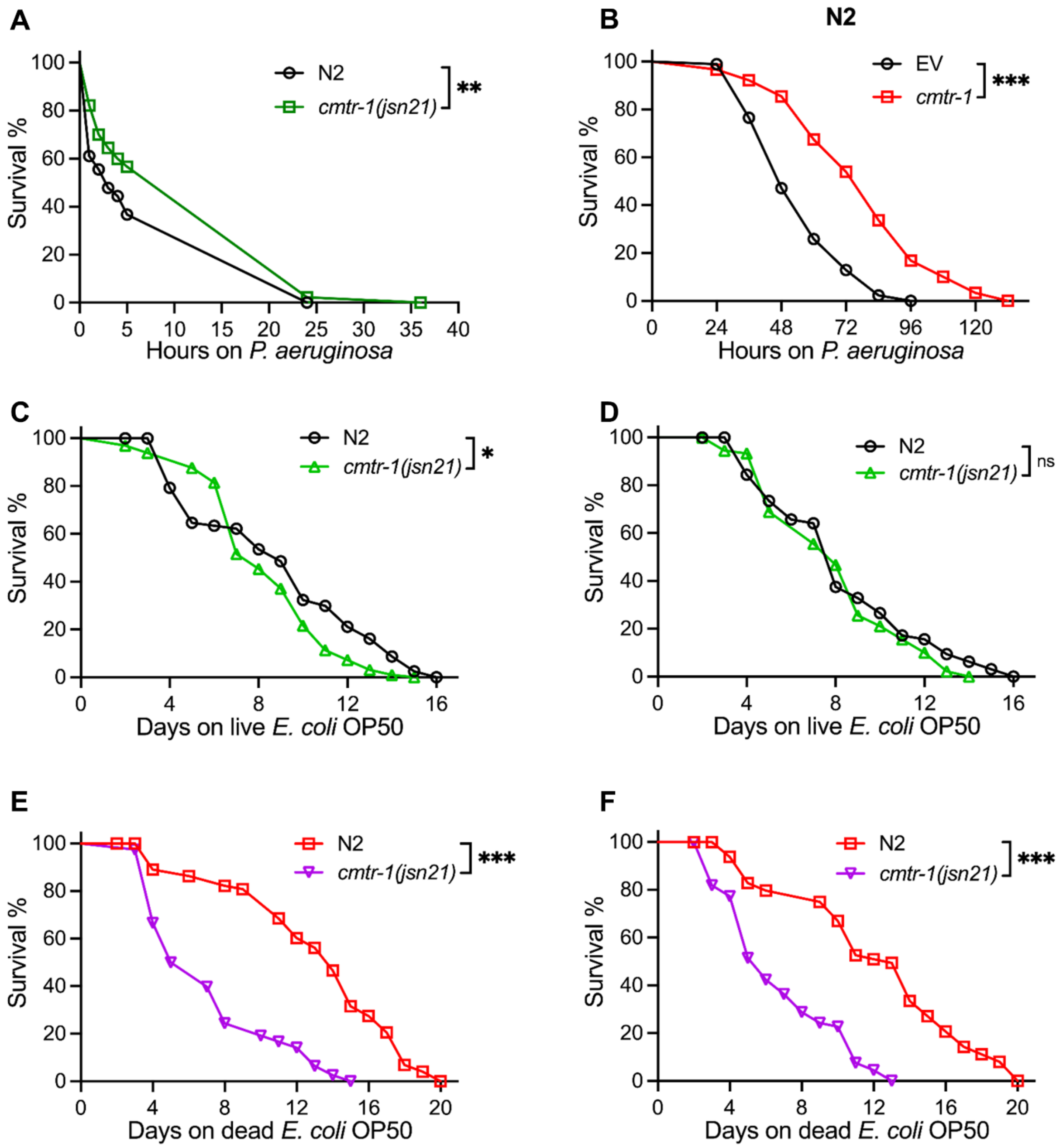


**Figure S2: Loss of *cmtr-1* enhances resistance to *P. aeruginosa* but does not extend lifespan**

(A) Representative survival plots of N2 and *cmtr-1(jsn21)* worms on *P. aeruginosa* PA14 at 25°C under fast-killing assay conditions. **$p < 0.01$ for the mutant as compared to the control worms ($n = 90$ per condition).

(B) Representative survival plots of N2 worms on *P. aeruginosa* PA14 at 25°C under slow-killing assay conditions after treatment with the empty vector (EV) control and *cmtr-1* RNAi. ***$p < 0.001$ for *cmtr-1* RNAi worms compared to the EV control worms ($n = 90$ per condition).

(C) Representative survival curves of N2 and *cmtr-1(jsn21)* worms fed on live *E. coli* OP50 in the absence of FUdR. *$p < 0.05$ for *cmtr-1(jsn21)* compared to N2 ($n = 92$ for N2 and 97 for *cmtr-1(jsn21)*).

(D) Representative survival curves of N2 and *cmtr-1(jsn21)* worms fed on live *E. coli* OP50 in the absence of FUdR. The difference between N2 and *cmtr-1(jsn21)* worms was not statistically significant ($n = 93$ for N2 and 100 for *cmtr-1(jsn21)*). Panels (C) and (D) are two independent biological replicates.

(E) Representative survival curves of N2 and *cmtr-1(jsn21)* worms fed on kanamycin-killed *E. coli* OP50 in the absence of FUdR. ***$p < 0.001$ for *cmtr-1(jsn21)* compared to N2 ($n = 82$ for N2 and 78 for *cmtr-1(jsn21)*).

(F) Representative survival curves of N2 and *cmtr-1(jsn21)* worms fed on kanamycin-killed *E. coli* OP50 in the absence of FUdR. ***$p < 0.001$ for *cmtr-1(jsn21)* compared to N2 ($n = 75$ for N2 and 69 for *cmtr-1(jsn21)*). Panels (E) and (F) are two independent biological replicates.

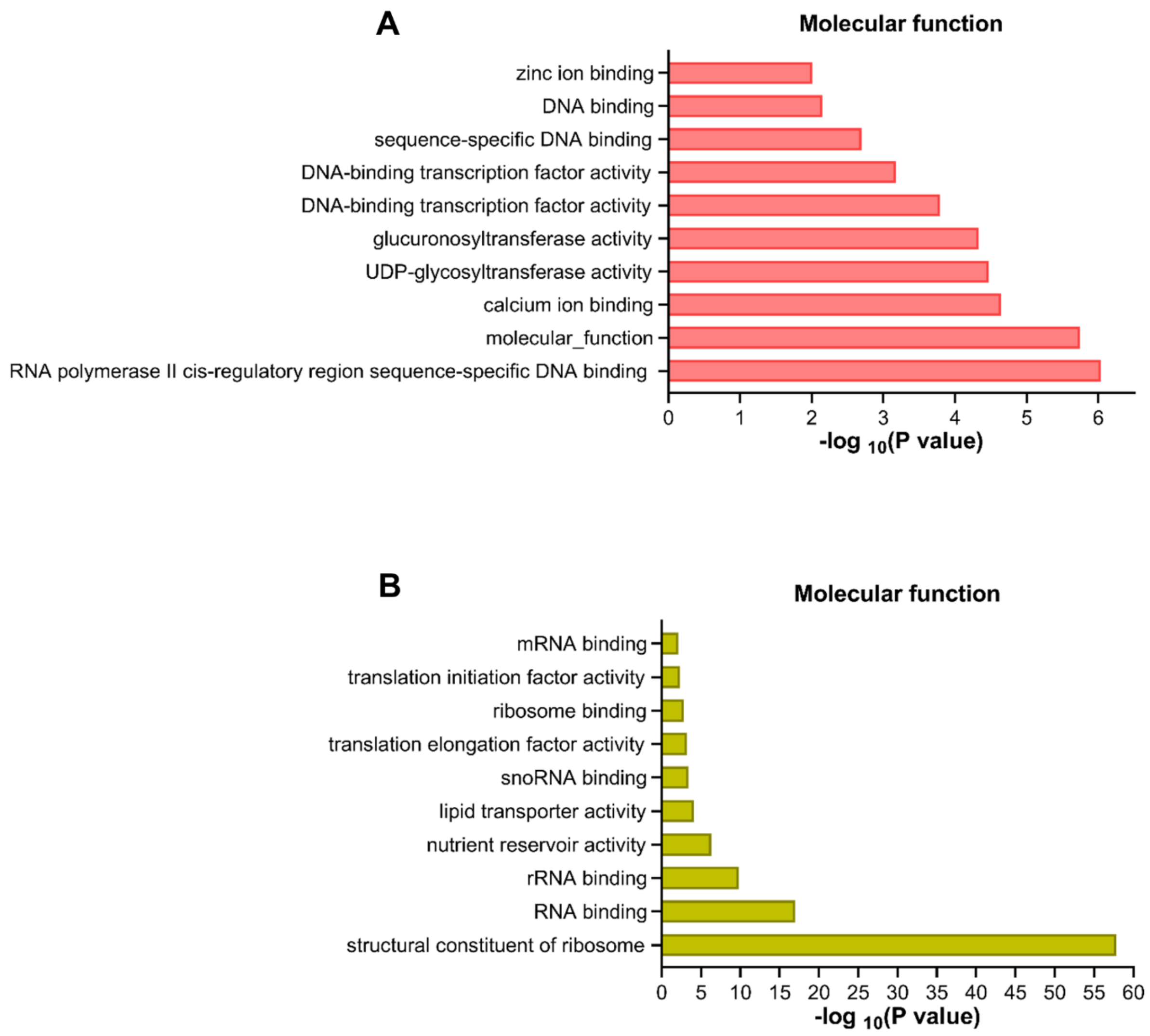


**Figure S3: *cmtr-1(jsn21)* mutants upregulate innate immune genes and downregulate translation-related genes**

(A)-(B) Gene Ontology (GO) enrichment analysis for molecular functions for upregulated (A) and downregulated (B) genes in *cmtr-1(jsn21);clec-60p::gfp* worms compared to control *clec-60p::gfp* worms grown on *E. coli* OP50.

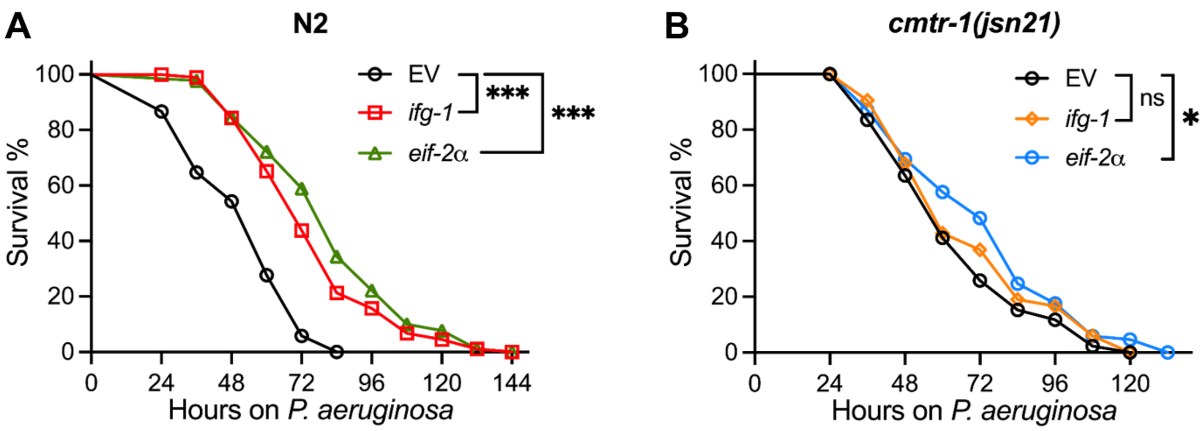


**Figure S4: Inhibition of translation initiation does not further enhance *cmtr-1(jsn21)* resistance to *P. aeruginosa* infection**

(A) Representative survival plots of N2 worms on *P. aeruginosa* PA14 at 25°C under slow-killing assay conditions after treatment with the empty vector (EV) control, *eif-2α*, and *ifg-1* RNAi. ***$p < 0.001$ for *eif-2α* and *ifg-1* RNAi-treated worms compared with EV control worms ($n = 90$ per condition).

(B) Representative survival plots of *cmtr-1(jsn21)* worms on *P. aeruginosa* PA14 at 25°C under slow-killing assay conditions after treatment with the EV control, *eif-2α*, and *ifg-1* RNAi. ***$p < 0.05$ and non-significant (ns) for *eif-2α* and *ifg-1* RNAi-treated worms, respectively, compared with EV control worms ($n = 90$ per condition).

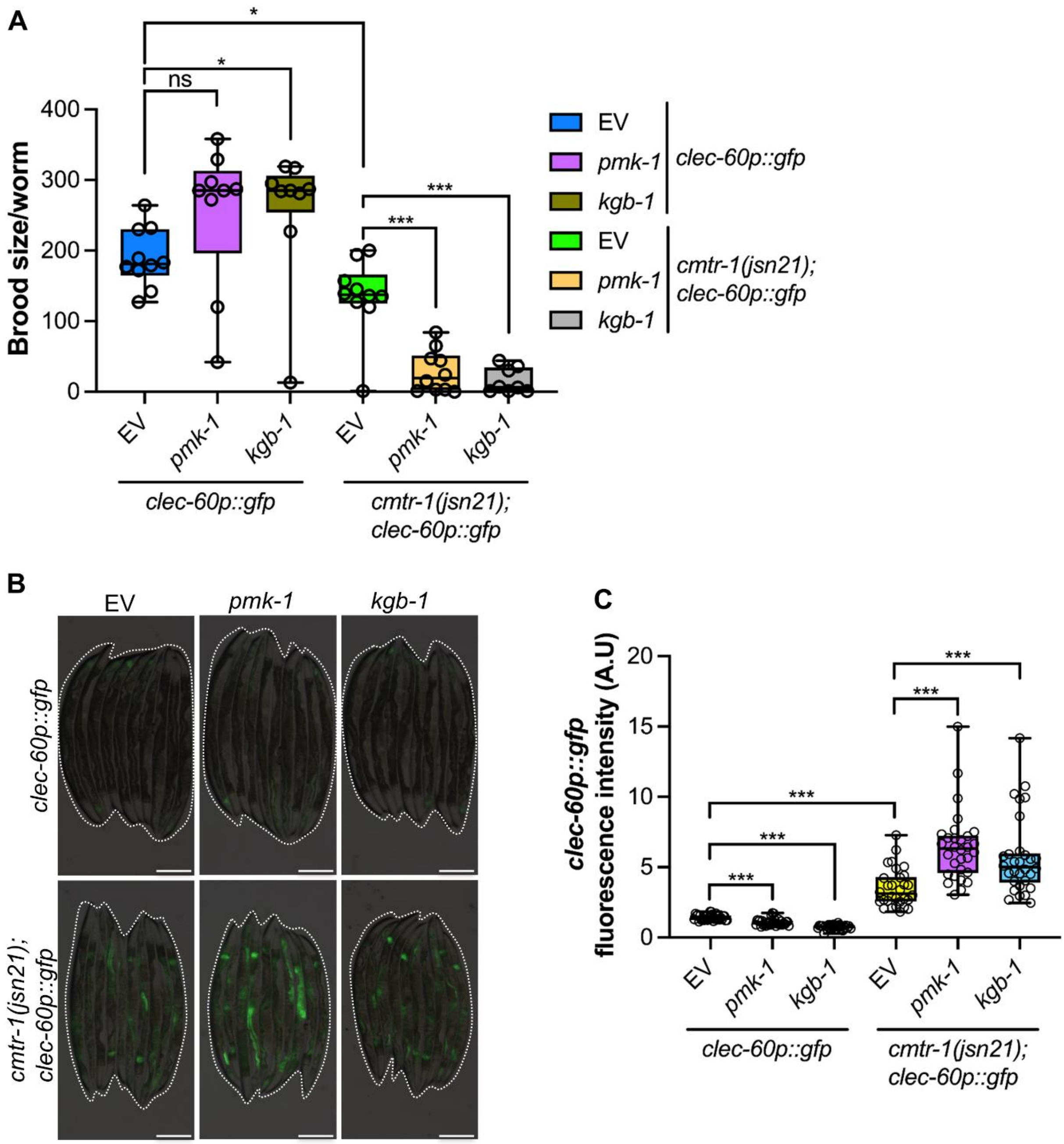


**Figure S5: Enhanced immunity in *cmtr-1(jsn21)* worms is independent of PMK-1 and KGB-1 pathways**

(A) Brood size of *clec-60p::gfp* and *cmtr-1(jsn21);clec-60p::gfp* worms grown on the empty vector (EV) control, *pmk-1*, and *kgb-1* RNAi. ***$p < 0.001$, *$p < 0.05$, and non-significant (ns) via t-test ($n$ = 8-10 worms each).

(B) Representative fluorescence images of *clec-60p::gfp* and *cmtr-1(jsn21);clec-60p::gfp* worms grown on EV control, *pmk-1*, and *kgb-1* RNAi. Dotted outlines indicate worm positions. Scale bar = 200 µm.

(C) Quantification of GFP levels of *clec-60p::gfp* and *cmtr-1(jsn21);clec-60p::gfp* worms grown on EV control, *pmk-1*, and *kgb-1* RNAi. ***$p < 0.001$ via t-test ($n$ = 30 worms each).

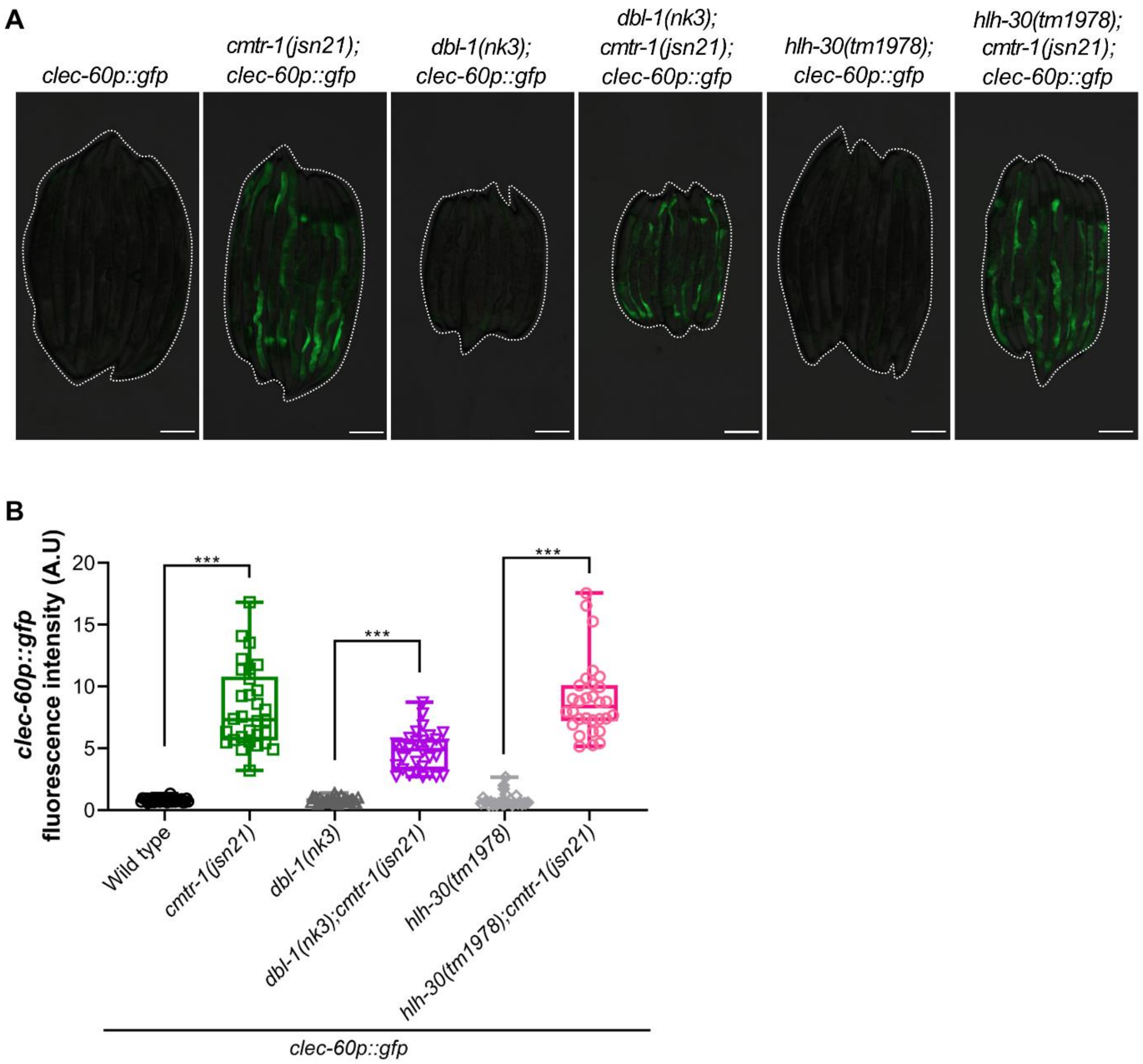


**Figure S6: Enhanced immunity in *cmtr-1(jsn21)* worms is independent of DBL-1 and HLH-30 pathways**

(A) Representative fluorescence images of *clec-60p::gfp*, *cmtr-1(jsn21);clec-60p::gfp*, *dbl-1(nk3);clec-60p::gfp*, *dbl-1(nk3);cmtr-1(jsn21);clec-60p::gfp*, *hlh-30(tm1978);clec-60p::gfp*, and *hlh-30(tm1978);cmtr-1(jsn21);clec-60p::gfp* worms. Dotted outlines indicate worm positions. Scale bar = 200 µm.

(B) Quantification of GFP levels of *clec-60p::gfp*, *cmtr-1(jsn21);clec-60p::gfp*, *dbl-1(nk3);clec-60p::gfp*, *dbl-1(nk3);cmtr-1(jsn21);clec-60p::gfp*, *hlh-30(tm1978);clec-60p::gfp*, and *hlh-30(tm1978);cmtr-1(jsn21);clec-60p::gfp* worms. ***$p < 0.001$ via t-test ($n$ = 29-30 worms each). The controls *clec-60p::gfp* and *cmtr-1(jsn21);clec-60p::gfp* are shared with Figure 4B.

**A**

| Rank | Motif | P-value | log P-pvalue | % of Targets | % of Background | STD(Bg STD) | Best Match/Details |
|---|---|---|---|---|---|---|---|
| 1 | CTCTTATC | 1e-21 | -5.040e+01 | 36.72% | 18.55% | 88.2bp (107.9bp) | elt-2/MA1701.2/Jaspar(0.900) More Information \| Similar Motifs Found |
| 2 | CTCCGCCCCT | 1e-13 | -3.017e+01 | 12.43% | 4.36% | 98.2bp (99.4bp) | POL003.1_GC-box/Jaspar(0.822) More Information \| Similar Motifs Found |
| 3 * | CATGGGCGATGA | 1e-11 | -2.742e+01 | 1.51% | 0.01% | 72.1bp (97.5bp) | egrh-1/MA2132.1/Jaspar(0.524) More Information \| Similar Motifs Found |
| 4 * | TTCATTTATG | 1e-11 | -2.693e+01 | 12.05% | 4.47% | 91.9bp (106.8bp) | LIN-39(Homeobox)/cElegans.L3-LIN39-ChIP-Seq(modEncode)/Homer(0.639) More Information \| Similar Motifs Found |
| 5 * | ATGATATCACTG | 1e-10 | -2.435e+01 | 2.07% | 0.09% | 98.8bp (93.9bp) | atf-7/MA1438.2/Jaspar(0.619) More Information \| Similar Motifs Found |
| 6 * | CACCAAAGAAAT | 1e-10 | -2.363e+01 | 2.07% | 0.10% | 106.2bp (99.6bp) | EGL-5(Homeobox)/cElegans-L3-EGL5-ChIP-Seq(modEncode)/Homer(0.544) More Information \| Similar Motifs Found |

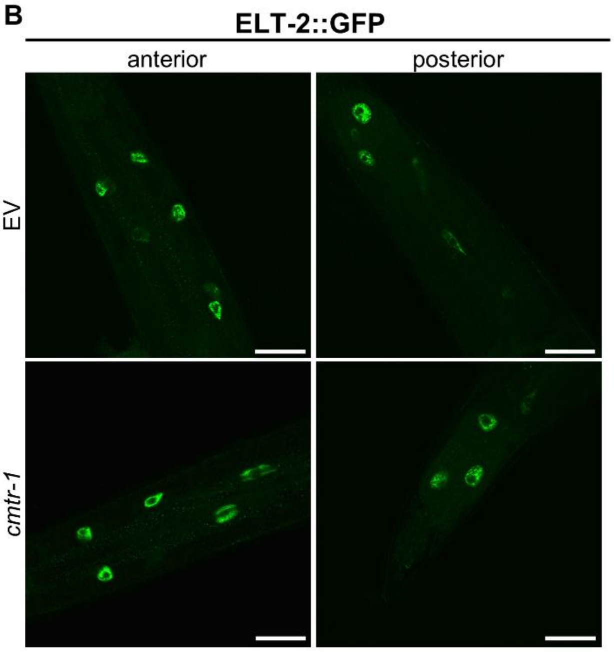


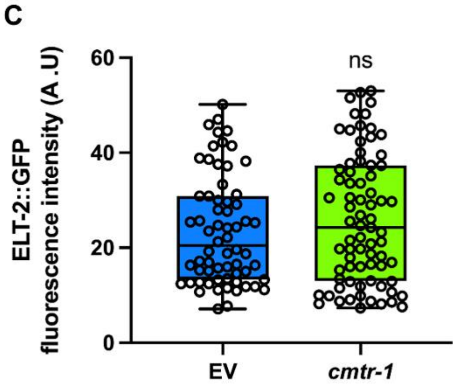


**Figure S7: The GATA transcription factor ELT-2 mediates immune activation downstream of *cmtr-1(jsn21)***

(A) List of the hits obtained for the most enriched binding motifs for the 1217 upregulated genes in *cmtr-1(jsn21)* worms, ranked in order from lowest to highest *p* values.

(B) Representative confocal fluorescence images of ELT-2::GFP worms grown on empty vector (EV) control and *cmtr-1* RNAi. Scale bar = 50 µm.

(B) Quantification of GFP levels of ELT-2::GFP worms grown on EV control and *cmtr-1* RNAi. Non-significant (ns) via t-test (*n* = 64 nuclei for EV and 75 nuclei for *cmtr-1* RNAi).

**Supplementary tables**

**Table S1:** Upregulated and downregulated genes in *cmtr-1(jsn21);clec-60p::gfp* worms versus *clec-60p::gfp* worms. Genes exhibiting at least two-fold change and *p*-value <0.01 were considered differentially expressed. A separate Excel file is provided.

**Table S2:** Primers used in the study. A separate Excel file is provided.

**Table S3:** Statistical analysis of survival curves from three independent experiments for each condition. A separate Excel file is provided.
